# Statistical power: implications for planning MEG studies

**DOI:** 10.1101/852202

**Authors:** Maximilien Chaumon, Aina Puce, Nathalie George

## Abstract

Statistical power is key for robust, replicable science. Here, we systematically explored how numbers of trials and subjects affect statistical power in MEG sensor-level data. More specifically, we simulated “experiments” using the MEG resting-state dataset of the Human Connectome Project (HCP). We divided the data in two conditions, injected a dipolar source at a known anatomical location in the “signal condition”, but not in the “noise condition”, and detected significant differences at sensor level with classical paired t-tests across subjects, using amplitude, squared amplitude, and global field power (GFP) measures. Group-level detectability of these simulated effects varied drastically with anatomical origin. We thus examined in detail which spatial properties of the sources affected detectability, looking specifically at the distance from closest sensor and orientation of the source, *and* at the variability of these parameters across subjects. In line with previous single-subject studies, we found that the most detectable effects originate from source locations that are closest to the sensors and oriented tangentially with respect to the head surface. In addition, cross-subject variability in orientation also affected group-level detectability, boosting detection in regions where this variability was small and hindering detection in regions where it was large. Incidentally, we observed a considerable covariation of source position, orientation, and their cross-subject variability in individual brain anatomical space, making it difficult to assess the impact of each of these variables independently of one another. We thus also performed simulations where we controlled spatial properties independently of individual anatomy. These additional simulations confirmed the strong impact of distance and orientation and further showed that orientation variability across subjects affects detectability, whereas position variability does not. Importantly, our study indicates that strict unequivocal recommendations as to the ideal number of trials and subjects for any experiment cannot be realistically provided for neurophysiological studies and should be adapted according to the brain regions under study.

## Introduction

Adequate statistical power is a requisite of robust, replicable science. An important variable affecting statistical power is sample size, and it has been shown that previous studies have been undermined by sample sizes that are too small (Button et al., 2013; Szucs & Ioannidis, 2017). Additionally, suboptimal scientific practices such as experimental designs and analysis approaches inappropriate to answering the posed scientific question have accentuated this problem (Gelman & Loken, 2013; Kerr, 1998; Kriegeskorte et al., 2009; Luck & Gaspelin, 2017). Overall, data gathering and analytical procedures that once were widely used are now identified as being flawed, while reporting procedures are codified (Keil et al., 2014; Pernet et al., 2020) and their endorsement is critically assessed (Clayson et al., 2019; Larson & Carbine, 2017). Emphasis is being put, on the one hand, on improving the robustness of statistical inference (Groppe, 2017; Kappenman & Keil, 2017; Kilner, 2013; Luck & Gaspelin, 2017; Simonsohn et al., 2013), and on the other hand, on the design, preparation, and documentation of carefully planned experiments (Chambers et al., 2015; Foster & Deardorff, 2017; Luck, 2005). The current study is at the crossroads of these two trends, aiming to aid researchers in cognitive, social and systems neuroscience in making principled decisions on how many trials and subjects to include in their experiments to achieve adequate levels of statistical power. More precisely, we highlight some important variables that one should pay attention to when considering how many repetitions of experimental conditions and how many subjects one should test to achieve robust statistical inference in an MEG experiment.

The question of knowing how many trials and subjects to include in an MEG or EEG experiment has to date been largely a matter of “know-how” or “rules of thumb” (Gross et al., 2013; Luck, 2012). Indeed, this topic has been discussed for decades without reaching any definitive conclusions (Duncan et al., 2009; Hari et al., 2018; Kane et al., 2017; Keil et al., 2014; Picton et al., 2000; Pivik et al., 1993). However, the above-mentioned concerns about power and reproducibility call for a systematic evaluation of variations in statistical power. This is particularly crucial in these current days where high-density MEG/EEG data are typically acquired. Which variables critically affect statistical power? Considering these, how can the necessary (and sufficient) number of trials and subjects be planned in advance? Recently, Boudewyn and colleagues (2018) took a first step at answering this question in a principled manner. They used EEG recordings from 40 participants to examine how the number of observations included in their analyses affected the probability of finding a significant effect in Event-Related Potential (ERP) measures. As expected, large effects (e.g., an error-related negativity, producing a 5-15 μV difference wave in the EEG) were detected at sensor level with fewer trials/participants than smaller effects were (e.g., a finer ~ 1 μV amplitude modulation in the lateralized readiness potential). We believe Boudewyn et al. (2018) to be the first EEG study that directly related the number of observations (trials and subjects) to statistical power. In another recent study, Baker and colleagues (2019) took this approach one step further, by acknowledging the difference between within-sample (i.e. inter-trial) and between-sample (i.e. between subjects) variability in a number of assessment modalities, including MEG and EEG. They introduced so-called “power contours”—plots that depict statistical power as a joint function of the number of trials and the number of subjects. These power contours reveal the level of statistical power reached for given trial and subject numbers. We illustrate the findings of the present study using this power contour plotting approach.

Here, we used MEG data from a large sample of subjects to reliably examine the behavior of statistical power measures. We simulated “experiments” with the open resting-state MEG dataset of the Human Connectome Project (HCP, Larson-Prior et al., 2013), varying the numbers of trials and subjects to assess the effects of within- and between-subject sample sizes, respectively. Importantly, we focused our analysis on the spatial properties of the neural source of the simulated effect. We sampled from a large cohort of subjects to approximate spatial variability across brains, but we did not attempt to model functional variability, in space or time. The only source of within-subject variability that we accounted for here is the level of physiological noise present in the data, which we approximated using resting-state HCP data from individual subjects.

This allowed us to simulate power in the context of real physiological background activity. We simulated effects by injecting dipolar sources of fixed amplitude at known anatomical locations in one half the data and not in the other half (Figure 1). We then detected the effects observed at sensor level with classical paired t-tests across subjects, corrected for multiple comparisons using a cluster-based approach. In an initial exploration, we used detailed individual source models and observed that detectability varies drastically according to the anatomical location of the source. Accordingly, we explored the spatial properties of the sources across the brain, focusing on their distance with respect to the closest sensor, their orientation with respect to the closest point on the sphere encompassing the subject’s head, and the cross-subject variability of these parameters. For this, we undertook two types of simulations. First, we examined changes of detectability across different anatomical areas, simulating sources in the real anatomy of individual subjects in our HCP dataset. In doing so, we observed that the four measured properties were impossible to properly disentangle due to anatomical constraints. Therefore, we ran a second set of simulations, where we imposed spatial properties of the sources independently of individual brain anatomy while using the same simulation strategy as before. In this way, we could assess separately the effect of each spatial property of the sources (position, orientation, and their cross-subject variability) on detectability.

**Figure 1.**
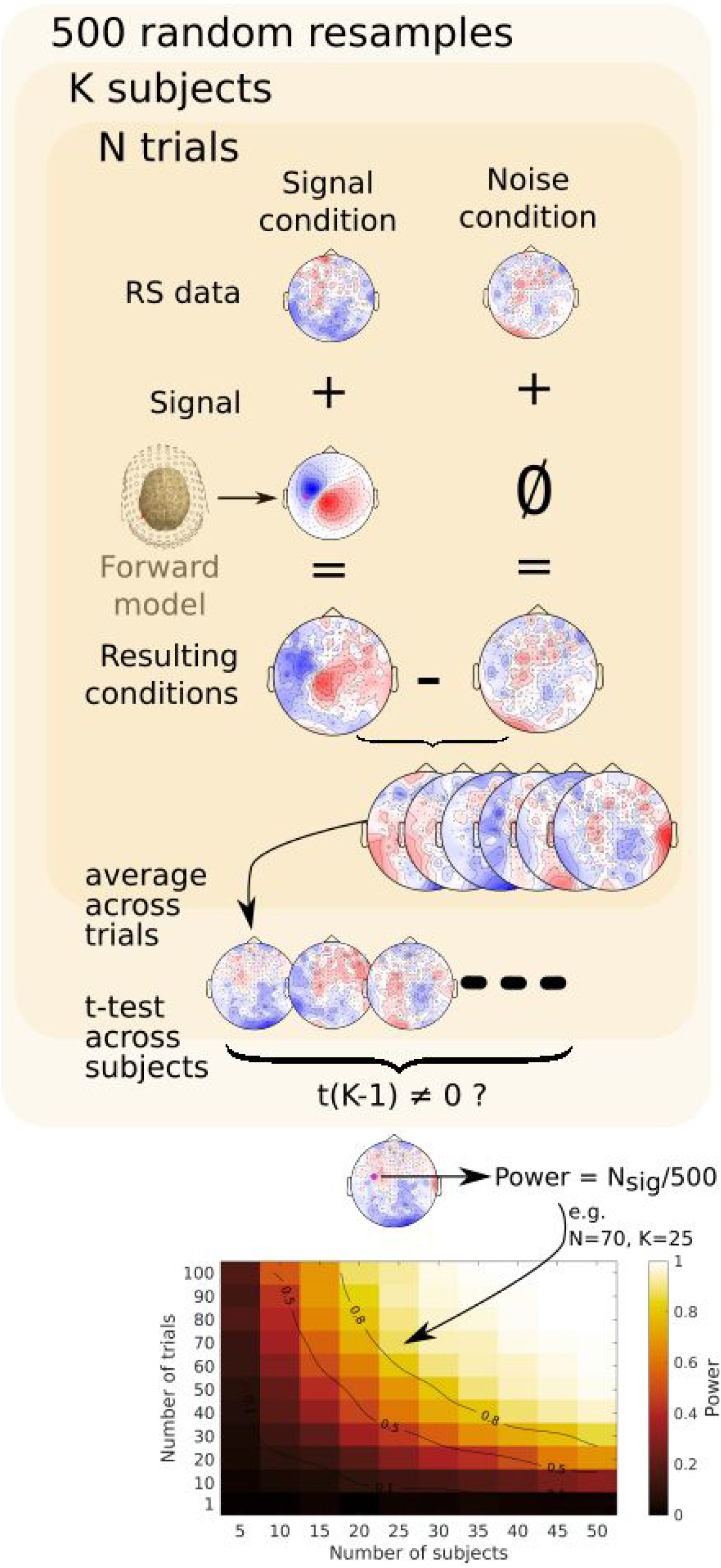
Schematic of simulation method. For a given “experiment” simulation with K subjects and N trials, K subjects were first selected at random. In each subject, N trials (that is, 50 time-point segments averaged along time) were randomly chosen and assigned to the two “conditions” from the Resting State (RS) data of the HCP. A signal created with the individual’s head model was added to the trials of the “signal” condition, whereas trials in the “noise” condition were unaltered. The data were averaged across the trials in each condition, the amplitude, squared amplitude, or global field power were computed, and the procedure was repeated for the K subjects. Significance was then tested across subjects by means of paired t-tests between conditions, corrected for multiple comparisons. The whole computation was repeated 500 times with different random samples of N subjects and K trials to yield an estimate of statistical power for this [K,N] pair, which was the number of significant permutation tests at the peak electrode (marked with a pink dot on the signal topography and on the statistics topography) divided by 500. This value was then color coded in the bottom power contour plot which represents power for every tested [K,N] pair. Contour lines in this and other power contour plots indicate power values (spline-interpolated for visualization purposes). These plots are used throughout the paper to illustrate the detectability of a given effect.

## Material & Methods

### Software

All data were processed using MATLAB (The Mathworks, Natick, MA, USA; Version 2018a) and the FieldTrip toolbox (Oostenveld et al., 2011, github release of October 25 2019). All analysis and visualization scripts and toolboxes, including the FieldTrip toolbox and megconnectome toolbox used, are available online at https://gitlab.com/icm-institute/meg/solid_meg.

### Data

#### *Input* data

We used MEG data from 89 subjects from the Human Connectome Project (HCP) open data repository, described in detail in Larson-Prior et al. (2013). Of the 100 recorded young adult subjects, 95 were included in the latest release (V3.0) of the MEG HCP data, 6 subjects with missing data in one or another of the components described below were discarded, resulting in the final number of 89 subjects for our analyses.

A whole series of preprocessing steps are already performed on the HCP data. For complete details, see the descriptions of the pre-processing pipelines for MEG (Larson-Prior et al., 2013), structural MRI (Glasser et al., 2013), as well as the online resources available at https://www.humanconnectome.org/software. We henceforth refer to these two preprocessing pipelines as the *HCP-MEG pipeline* and the *HCP-structural pipeline*. Note that these pipelines are part of different releases of the HCP data that were downloaded separately from https://db.humanconnectome.org. Subject identifier codes being consistent across these releases, we used them to merge the data from both pipelines.

Briefly, the MEG data were acquired at 2043.5101 Hz (bandpass filtering: DC-400 Hz) with a whole-head 248 magnetometer device (MAGNES 3600, 4D Neuroimaging, San Diego, CA). The HCP includes data from several activation tasks and three 6-minute resting state periods (eyes open, fixating). We used the latter dataset in this study. MEG fiducials were obtained with a Polhemus system for coregistration to structural MRI data. Each subject was scanned with a high-resolution structural MRI on a 3T Siemens Skyra MRI scanner (Van Essen et al., 2012). Within the HCP-MEG pipeline, the structural MRI was used to create a single-shell volume conduction model using FieldTrip. Furthermore, to define anatomical labels in our study, we used the output of the HCP-structural pipeline, where a high-definition segmentation and anatomical labelling (Destrieux Atlas) of the cortical mantle was performed using Freesurfer (Version 5.2, Fischl, 2012).

All of the material described above is available from the HCP database (individual MEG sensor space time courses, magnetometer definition, individual source space and individual head model) and formed the input data for our study.

#### Head model, source space, and forward model

We used the head models as provided for each subject by the HCP-MEG pipeline. We used the 4D Neuroimaging/BTi coordinate system to identify source positions within these models throughout the paper^1^ (with mm units for clarity). We created a leadfield matrix with FieldTrip using the magnetometer description and the provided head models, and generated the sensor signals by projecting sources (see below) with a 10 nA.m amplitude through the leadfield to the sensors.

We used two different types of source models, where the source dipoles were either constrained to be orthogonal to the individual cortical mantle, or free from any anatomical constraint, as explained below.

#### Source models constrained by anatomy

We performed a first set of simulations constrained by individual subjects’ anatomy, where the signal to be added to the resting-state data was generated from one source (i.e. one vertex) of the cortical mesh provided by the HCP. This source was oriented orthogonally to the cortical mesh. Across participants, we identified sources of similar areal origin by means of their index in the cortical mesh provided by the HCP. It is worth mentioning that we do not imply that these indices map strictly homologous portions of cortex across subjects. The mesh topology is, however, identical in all 89 subjects (all 64,984 vertices, spaced on average by 1.5 mm, are connected in the same way in every subject), and it is used in practice to make cross-subject source-level comparisons and averaging (Fischl, 2012). No within- or between-subject variability other than the one already present in the HCP data was added. Between-subject variability thus occurred because of variations in position and orientation of the source vertices across subjects.

For high-resolution rendering of the cortical surface in figures, we used the high-definition segmentation with 163,842 vertices, spaced on average by 0.5 mm, per hemisphere found in the HCP-structural pipeline (Glasser et al., 2013). All segmentations used in this study are linearly coregistered to MNI coordinates (same origin, scale, and orientation), but not “warped” (i.e. non-linearly transformed to closely map to the MNI template brain) as is often performed for mapping individual source reconstructions across participants in MEG. This non-linear mapping would have been inappropriate here since we were interested in between-subject anatomical variability.

All our brain anatomy-constrained simulations used an equivalent current dipole of 10 nA.m as the source of difference between the two conditions. Thus, we did not vary signal amplitude in this study. In a recent study, Murakami and Okada (2015)showed that current density due to local neuronal currents, expressed as equivalent current dipole per unit of surface (in nA.m/mm^2^), is remarkably constant throughout a range of vertebrate species. They argued that a value of current dipole moment density of 0.16-0.77 nA.m/mm^2^ may serve as an effective physiological constraint for human neocortical MEG/EEG sources. As an indication, according to these estimates, the current dipole value of 10 nA.m that we chose in this study could correspond to an activation surface ranging between 13 and 62 mm^2^. However, complexities arise due to the spatial extent of the sources (e.g. changing orientations around convexities of the cortical mantle). Accounting for these complexities is beyond the scope of this paper, but it has been tackled in previous studies (Ahlfors, Han, Lin, et al., 2010; Fuchs et al., 2017).

#### Source models unconstrained by anatomy

As will become clear, measures of source properties that affect detectability tend to covary across the brain in ways that are not readily predictable (Supplementary Figure 2). We thus also used a simulation approach where dipoles were placed independently of cortical anatomy. For these simulations, we took an *initial source* located within the precentral region (x=−12, y=33, z=70; source position illustrated in Figure 8), oriented it normally to the mean cortical surface at that location, and systematically varied the spatial source properties starting from that location (see “Source properties description”, below).

All other parameters of the simulations were kept identical to those of the anatomically constrained simulations described above, except that the amplitude of the injected sources was reduced to 5 nA.m to avoid strong saturation of the power contour plots due to the very weak between-subject variability in simulations where individual parameters were held constant (see details below).

### Simulations

In all of our simulations, we used the same procedure based on a Monte Carlo resampling strategy. The Monte Carlo procedure uses repeated random selection from a data sample to approximate a property of the general population, that is, here, the probability of finding a statistically significant effect in the general population (a.k.a. statistical power).

As illustrated in Figure 1, for each simulation with a given number of subjects and trials, we first randomly chose the required number of subjects from the HCP database, sampling without replacement. For each subject, we then randomly selected our “trials”. Each trial consisted of 25-ms (50 samples) time segments of the continuous resting state data, at least 2 s apart from each other. We then split these trials randomly in two sets of equal size, added signal (according to the procedure explained above) in one set and averaged the data across time points and trials separately for each set. So, our two conditions consisted of “signal” and “noise” trials. We then ran a paired t-test of the difference between the two sets across subjects at each sensor and noted significance (p<0.05, uncorrected). To summarize the whole MEG helmet with one value, we used the state-of-the-art spatial cluster-mass based correction for multiple comparisons (Maris & Oostenveld, 2007; 1000 permutations, cluster and significance thresholds both at p<0.05) and considered a given comparison significant only when the peak sensor (i.e. the sensor at which the absolute value of the projection of the source signal peaked) was included in a significant cluster. We repeated this procedure 500 times for each trial-by-subject number pair and noted the number of times where the comparison was significant among these 500 simulations. This so-called Monte Carlo statistical power estimate approximates the probability of finding a significant effect if we were to run an experiment with the given parameters (number of subjects, trials and signal properties) in the general population.

In an initial simulation, we assessed overall variations in statistical power for simulated sources throughout the brain with the “rule of thumb” number of subjects and trials (25 subjects, 50 trials per condition for each subject). This is reported in the initial results section, “*Overall variations of statistical power throughout the brain*”.

In subsequent simulations, we ran Monte Carlo simulations for all combinations of trial and subject numbers, from a single trial to 100 trials (in steps of 10) in each condition and with 5 to 50 subjects (in steps of 5), in order to assess the impact of different source properties on statistical power as described below.

For all analyses, we ran paired t-tests on amplitude differences, as well as on differences in squared amplitudes (a.k.a. signal power) and on the difference in standard deviation of amplitudes across sensors (a.k.a. global field power, or GFP, or global mean field power, GMFP, Lehmann & Skrandies, 1980), because these are common measures applied to data in sensor-level analyses.

### Source properties description

We used the simulation approach described above to evaluate how variations in the spatial properties of the source dipole affect signal detection at sensor level. We examined what we call first- and second-level properties of the sources. First-level properties refer to the position and orientation parameters of the sources. We defined position as the distance between the source and the closest sensor, and orientation as the orientation of the source with respect to the closest point on a sphere fitted to the subjects’ head. Second-level properties refer to the *variability* in first-level properties across subjects. We examined separately the first-level properties (position and orientation) and second-level properties (cross-subject variability in position and in orientation), in four sets of simulations. Each property was first examined in simulations of sources using individual brain anatomy, then manipulated in simulations unconstrained by anatomy.

#### First-level properties: position and orientation of the source

The following two measures allowed us to explore the extent to which effect detectability depends on the distance of the source with respect to the sensor array and on its orientation with respect to the sphere encompassing the subjects’ head.

#### Distance to the closest sensor

Distance from the source to the detector is a major determinant of the measured signal amplitude. In simulations constrained by individual brain anatomy, the distance to the closest sensor for a given source was measured on the individual anatomy in each subject. In simulations not constrained by anatomy, we imposed this distance by shifting the position of the initial source (described in the “*Models unconstrained by anatomy*” section above) by a target distance inwards or outwards, along the line passing through the sensor closest to the initial source (labeled A44 in the original data) and the origin of the coordinate system, so that the distance to the closest sensor ranged from 40 to 120 mm. A source property more commonly assessed when studying source detectability is source depth in the head (i.e. distance from the source to the scalp; Hämäläinen et al., 1993). We chose distance to sensor instead because it is ultimately the one that matters for signal detection, and to account for uncontrolled head-size variability in depth.

#### Orientation angle with respect to the head surface

Dipolar sources oriented radially within a spherical volume conductor produce no net magnetic field outside of this volume. Even if the sphere is a gross approximation of the subjects’ head volume, the orientation angle of brain sources with respect to the sphere encompassing the subjects’ head is often used as a reference to predict the detectability of a source. Here, we computed the orientation angle of brain sources with respect to the closest point on a sphere of 10 cm radius fitted to the shape of the single-shell head model provided by the HCP MEG pipeline. The orientation angle, expressed in degrees, was computed as the arccosine of the product of the orientation vector of the source with the orientation vector normal to the closest point on the sphere. It ranged from 0° for sources pointing directly towards the sphere surface (i.e. radial sources pointing outward the head surface), through 90° for sources pointing orthogonal to the sphere surface (i.e. tangential sources), up to 180° for sources pointing directly away from the sphere surface (i.e. radial inward-directed sources).

The source orientation angle was normal to the cortical surface in simulations using individual brain anatomy. It was manipulated parametrically in simulations not constrained by anatomy, by rotating the initial source dipole around the y-axis of the coordinate system (that is, the axis passing through both ears). This axis approximates, for the initial source used in these simulations, the rotation from a radially oriented source on the precentral gyrus to a tangentially oriented source such as the sources located on the wall of the nearby central sulcus.

#### Second-level properties: variability of the sources across subjects

Cross-subject variability in source position and orientation are thought to decrease detectability at the group level, and were thus examined here.

#### Position variability

Position variability was defined as the standard deviation of source position across subjects. In simulations not constrained by individual anatomy, we sampled dipole locations at random from an uncorrelated trivariate (x, y, z) normal distribution with a standard deviation varying from 0 to 10 mm. Put in another way, in these simulations, the dipoles were varied randomly across subjects so that approximately 95% of the dipoles were encompassed within a sphere with a radius about twice the given standard deviation.

#### Orientation angle variability

To measure orientation angle variability, we computed the length of the average orientation vector across subjects for each source location and we took the log of the inverse of this value, so that smaller values (lower bound of 0) reflect smaller variability and higher values (upper bound of infinity) reflect higher variability. This measure is unitless. In simulations not constrained by anatomy, orientation variability was imposed by adding an uncorrelated random bivariate (azimuth and elevation) normally distributed angle to the individual sources with a standard deviation varying from 0 to 180°.

## Results

### Overall variations of statistical power throughout the brain

Before getting into the description of the explored spatial properties, here we describe the overall variations of statistical power for detecting sources throughout the brain. We considered three types of signal measures that are commonly used in sensor-level analyses: amplitude, squared amplitude (or power), and standard deviation of amplitude across sensors (or global field power, GFP). In Figure 2, we present statistical power as computed for these three measures at sensor level. We calculated the average statistical power for detecting a source at each possible cortical location for a highly detailed source model following individual brain anatomy, in a simulated dataset of 25 human subjects and 50 trials per condition. Such subject and trial sample size is quite typical in cognitive and social neuroscience studies.

**Figure 2.**
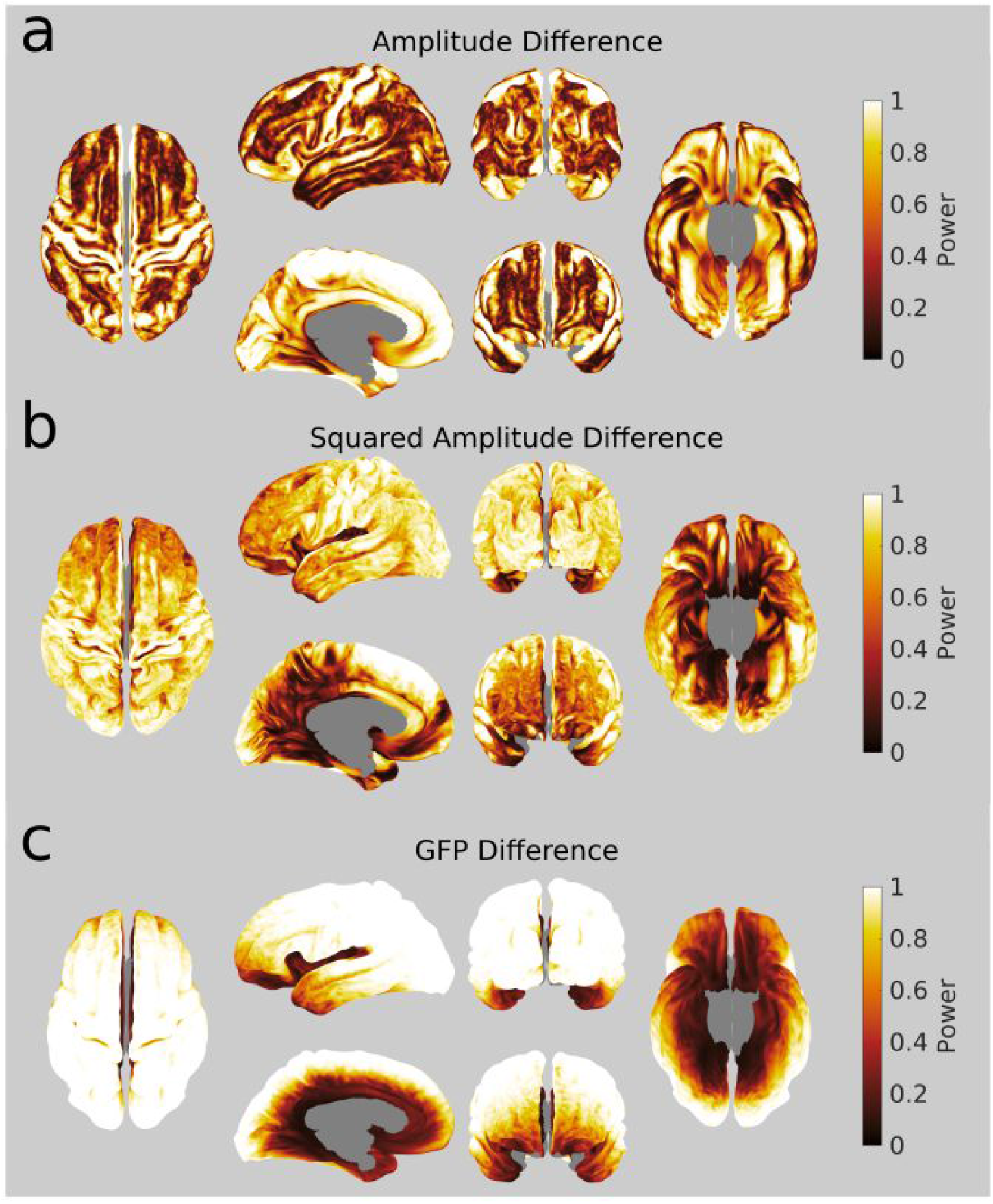
Variation in estimated statistical power across the cortical surface with three different signal measures for simulated evoked activity. Statistical power estimated from three different measures for detecting (at MEG sensor level) the activity generated by a single dipole placed in turn at every possible cortical position. At each position, we simulated experiments with 25 subjects with 50 trials each. Images of superior, left lateral, left medial, anterior, posterior, and inferior brain views are depicted. **a**. Amplitude difference. Note the relatively high statistical power values within major sulci and on the medial wall. **b**. Squared amplitude difference. Note the relatively lower statistical power in medial and inferior regions **c**. GFP difference. Note the relatively high statistical power at superficial neocortical sources, and low statistical power in deep sources. The paired differences between these three maps are shown in Supplementary Figure 1.

Figure 2 presents the results of this initial analysis. Note that the absolute statistical power values *per se* are somewhat arbitrary, because we simulated the effect of dipoles with an arbitrary, constant amplitude. What is important is the striking heterogeneity of statistical power across the brain that this figure reveals and the complex dependency of statistical power on the type of measure considered. Some of this heterogeneity may appear trivial with regard to what is known on MEG signal sensitivity to source orientation and depth. For instance, sources at gyral crests tend to be detected with low statistical power, probably because they are radially oriented. In contrast, sources on the walls of the central sulcus and on the medial surface of the frontal lobe—which are tangentially oriented—are detected with high power. However, some other observations may appear to be far less intuitive. For instance, the ventral surface of the temporal lobes shows relatively high power in comparison to the lateral surfaces, even though it is further away from the sensors. In addition, the results obtained using different measures of the sensor level signal (Figure 2b, and c) also revealed that the computation used to quantify activity affects statistical power. This observed heterogeneity of statistical power across the brain and across measures calls for a better understanding of the influence of source properties on the sensor-level detectability of generated effects. For this, in the following sections, we examined the first- and second-level spatial properties of the sources, i.e. the spatial parameters defining source position and orientation, and their variability across subjects.

### First level properties: Distance and orientation of sources

#### Distance

Figure 3a depicts the average distance to the closest sensor for every source location on the cortical surface, plotted on the average brain of the 89 subjects in the HCP database. The histogram of this distance across all vertices is plotted in Figure 3b. The distance varies from about 4 cm in cortical regions closest to the sensor array (central sulcus, occipital cortex) to about 10 cm for the deepest structures (cortex near the hippocampus, amygdala, and basal forebrain).

**Figure 3.**
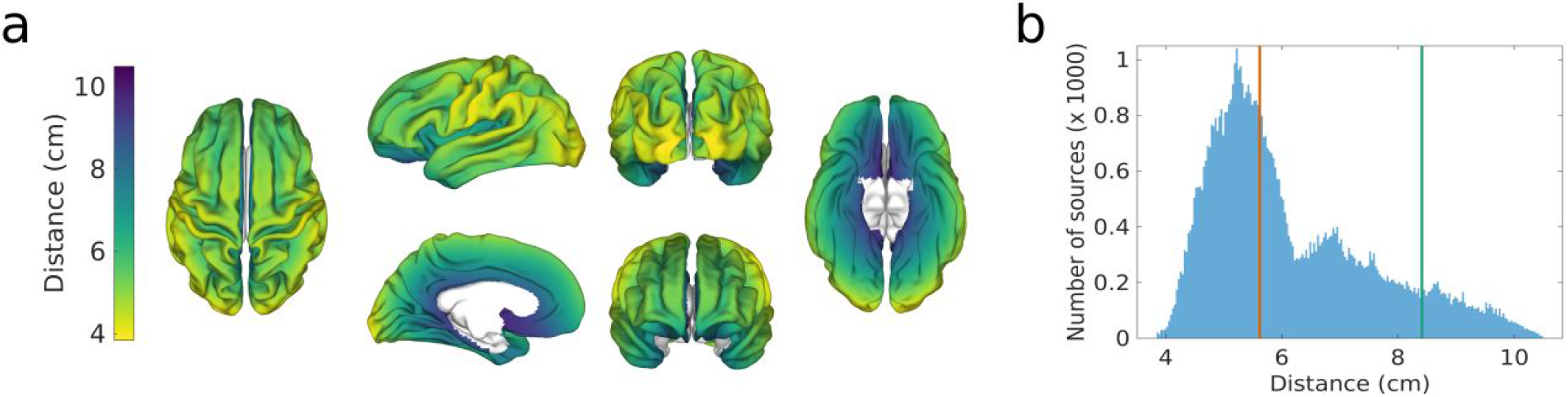
Mean distance to the closest sensor across subjects. Data calculated from 89 subjects in the HCP MEG dataset and presented on the average brain. **a. Mean distance to the closest sensor**, mapped on average brain views (from left to right and top to bottom: superior, left lateral, left medial, posterior, anterior, and inferior views). Greater distances appear in darker color. **b. Histogram of the distribution of the mean distance to closest sensor across all cortical vertices.** The red and green vertical lines indicate the distance to the closest sensor of the two dipoles that we selected to illustrate the effect of this parameter in Supplementary Figures 3 and 4, respectively.

The effect of distance to sensors on statistical power is, as expected, a decrease in power with increasing distance from the sensors. For brevity, we illustrate this effect in detail in Supplementary Results (see Supplementary Figures 3, 4, and 5). In brief, the distance to the sensors affects detectability drastically. A source with a given amplitude and orientation that is readily detected with very few trials and subjects when placed at a cortical location close to the sensors is virtually impossible to detect if it occurs near the center of the head. However, in such a situation, changing the measure used to test for differences can have a strong effect on statistical power.

It may be noted that deep sources (that is, sources with large distance to sensors) usually have a diffuse distribution of signal across multiple sensors. Therefore one could wonder if averaging signal over multiple sensors would augment deep source detectability. We found that this was not the case. To test this, we replicated our analysis of the effect of distance to sensor for sources unconstrained by brain anatomy (illustrated in Supplementary Figure 5), averaging signal measurements over 2, 5, or 10 sensors around the maximal sensor. No change in statistical power could be observed.

#### Orientation

Figure 4a shows the average source orientation relative to the closest point on a sphere encompassing the head of the subjects for every location on the cortical surface, plotted on the average brain of the 89 subjects in the HCP data. Most sulcal walls, a large part of the inferior temporal cortex and of the medial wall are close to tangential orientation (0°), whereas the crest of gyri (e.g. superior frontal gyrus) and the depth of sulci (e.g. the insular cortex) are closer to radial orientation (90°). The histogram of source orientations across all vertices is plotted in Figure 4b. It is evident that the distribution is skewed towards values below 90°, reflecting the overall convex topology of the cortical mantle.

**Figure 4.**
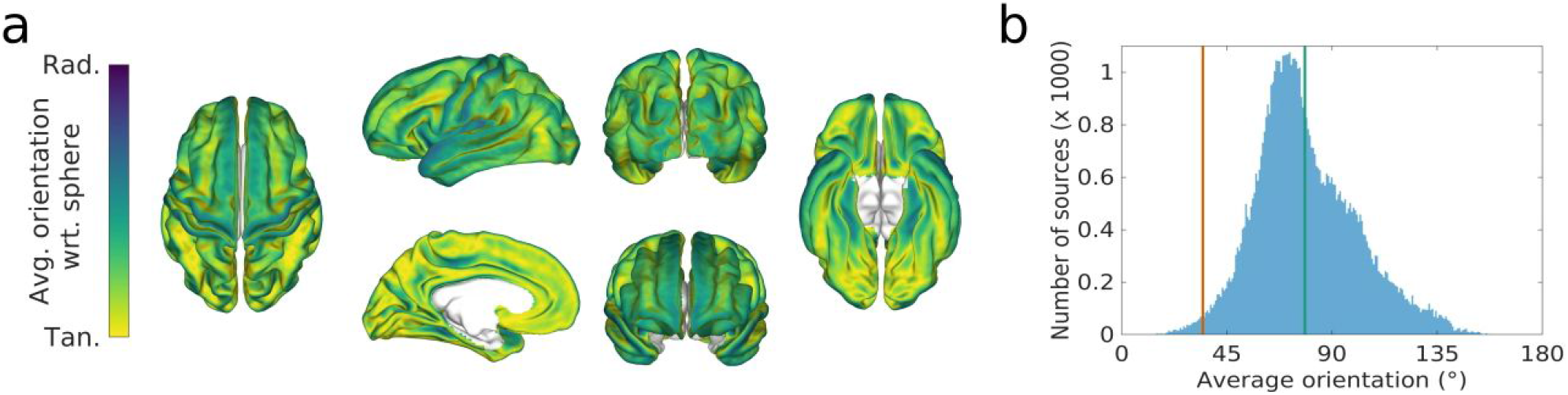
Average dipole orientation with respect to the closest point on the sphere encompassing the head across subjects. Data from 89 subjects in the HCP data plotted on the average brain. **a. Average orientation relative to the closest point on the sphere**. The figure shows the mean orientation mapped on superior, left lateral, left medial, posterior, anterior, and inferior views of the brain (from left to right and top to bottom). For illustration clarity, here, we mirrored the computed angle around the tangential axis (90°). In other words, we discarded information on whether the dipoles pointed towards or away from the sphere surface and only considered orientation (numerically, the values displayed are the inverse cosine of the absolute value of the product of interest). Legend on color bar: Rad. = radial (0 or 180°), Tan.= tangential (90°), Avg. = average, wrt. = with respect to. **b. Histogram of the distribution of mean orientation across all cortical vertices**. Contrary to panel a, values spread the full half circle. The red and green vertical lines indicate the orientation of the two dipoles that we selected to illustrate the effect of this parameter in Supplementary Figures 6 and 7, respectively.

The effects of source orientation with respect to the sphere encompassing the subject’s head are, as expected, that radial sources are very difficult to detect. For brevity, we show these effects in detail in Supplementary Results (see Supplementary Figures 6, 7, 8). In brief, source orientation affects detectability drastically, especially when sources are pointing exactly towards (or away from) the head surface (i.e. radially). However, this loss of power is recovered rapidly as soon as the orientation of the sources shifts away from the radial orientation. Moreover, the effect of orientation on statistical power is greatly affected by the type of signal measure applied to the data before testing. Differences between GFP measures, and to a lesser extent differences of squared amplitudes, can increase statistical power for detecting sensor-level effect of radial sources, especially when superficial sources are considered (see Figure 2, and Supplementary Figure 1).

Taken together, the results above allowed us to explore the effect of the position and orientation of dipolar sources on statistical power for detecting effects at the sensor level. Quite predictably, we have seen that these first-level properties have a strong impact on signal detectability. We now turn to our explorations of second-level spatial properties, and their effect on group-level source detectability, where the observable effects can at times be somewhat unexpected.

### Second-level properties: Position and orientation variability across subjects

#### Position variability across subjects

Figure 5a shows the average cross-subject standard deviation (across the three cartesian dimensions) in *position* for every source location on the cortical surface. The variability histogram across all vertices is plotted in Figure 5b. Overall, values ranged from 3.4 to 8.1 mm, with maximal values occurring in areas where cortical folding is more variable across subjects, such as in occipital cortex, and minimal values occurring in anterior medio-ventral region and in insular cortex. Dipoles placed in the most variable regions could be up to 10 mm away from each other, whereas dipoles placed, for example, in the insular cortex, are on average within 5 mm from each other.

**Figure 5.**
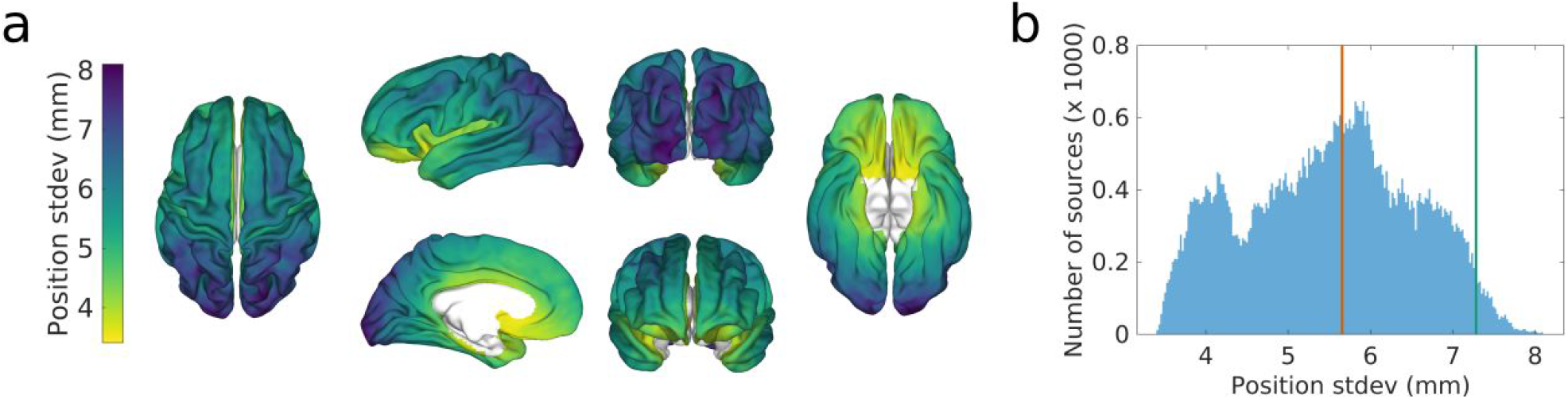
**Average cross-subject variability in position** shown on the average brain of 89 subjects in the HCP dataset. **a. The cross-subject variability in position** is the standard deviation of position of the sources across subjects, shown on superior, left lateral, left medial, posterior, anterior, and inferior views of the brain (from left to right and top to bottom). **b. Histogram of the distribution of average standard deviations across all cortical vertices.** The red and green vertical lines indicate the orientation of the two dipoles that we selected to illustrate the effect of this parameter in Figures 6 and 7, respectively.

Position variability is expected to have an impact on detectability because dipoles at different locations, even if they share the same orientation, project a magnetic field that is topographically different. However, changes in position will rarely change the polarity of the magnetic field at a given sensor, and the similarity in the topographies created by two dipoles may override their differences. So, it is hard to predict the extent to which variability in position will affect detectability at sensor level. To address this question, we first selected two posterior locations in the individual brains of the HCP subjects, one with relatively low spatial variability in fusiform gyrus (x=−39, y=33, z=5), and the second one with large variability in superior occipital cortex (x=−69, y=12, z=48).

For the fusiform gyrus source, the dipole was contained within a narrow +/− 10 mm region in the horizontal plane (x-y plane in Figure 6a) and within +/− 15 mm in the vertical plane (x-z plane in Figure 6a), across all subjects (see also Figure 6b). The orientation of this dipole was relatively consistent across subjects (Figure 6c). This source resulted in a signal with a strong amplitude at sensor level (Figure 6d), despite being at a considerable distance from sensors, and it was thus detected with a high statistical power across subjects, with 80% power observed with as little as 15 subjects with 60 trials each, or only 20 trials in 35 or more subjects (Figure 6e). When using the difference of squared amplitudes at sensors, or of GFP, to compare conditions statistical power was generally decreased.

**Figure 6.**
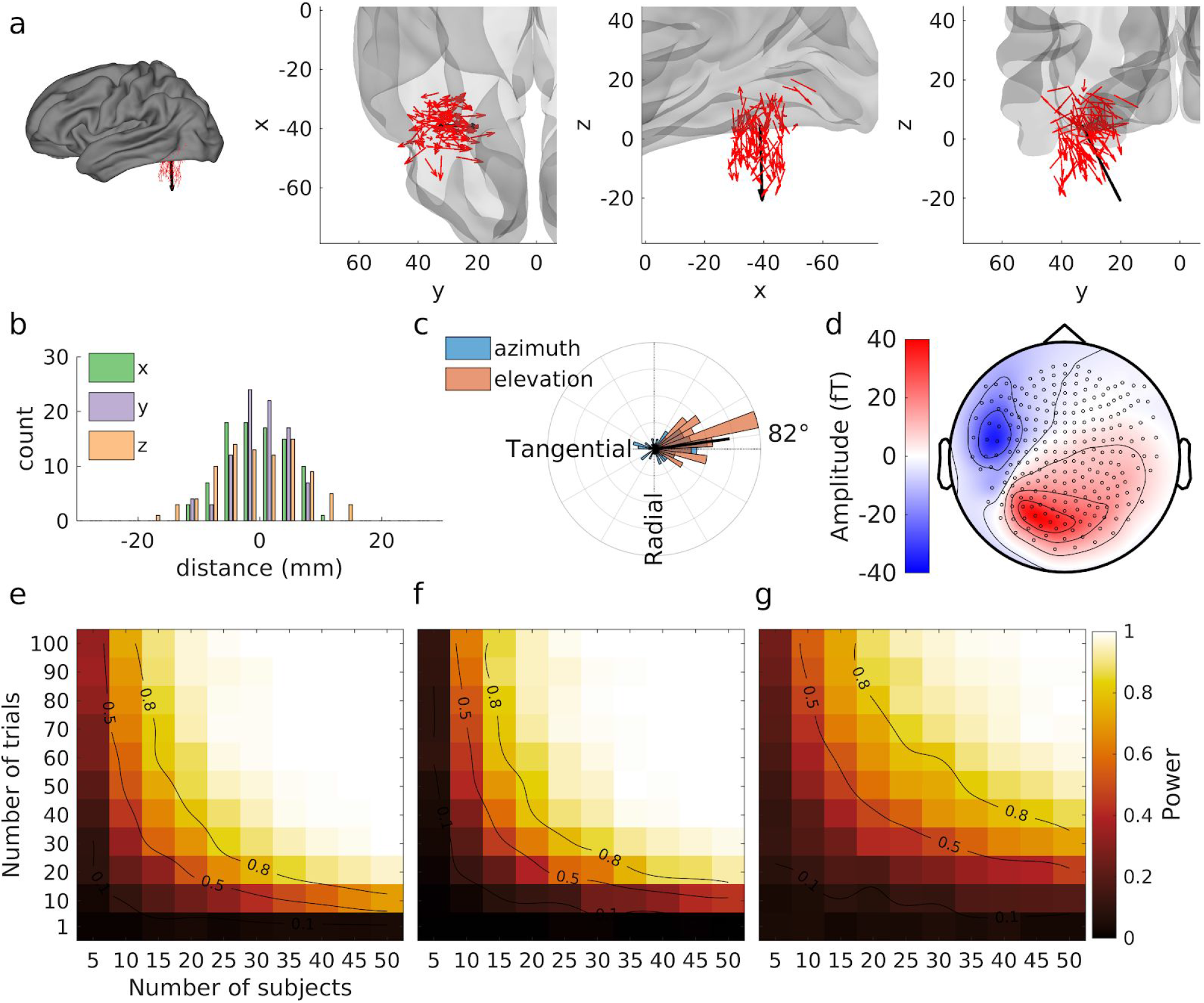
Detecting sensor-level effects for a source with relatively low cross-subject position variability in the fusiform gyrus. **a**. positions of individual dipoles in 89 subjects (red arrows) are represented on the average brain in a superior overall view, and in zoomed superior, lateral, and posterior partial brain views. The bold black arrow in each panel shows the resultant vector or average dipole across 89 subjects. **b. Histogram of the distance of each dipole to the mean position of all dipoles in the three cartesian dimensions. c. Polar histogram of the orientations of the individual dipoles relative to the orientation of the closest point on the sphere encompassing the subjects’ head**. Azimuth and elevation are referenced to the closest point on the sphere surface, so an orientation orthogonal to that of the sphere encompassing the subjects’ head is shown as tangential (90°), and the colinear orientation is shown as radial (0°). The black thick line coming outwards from the center of the plot represents the orientation and length of the resultant vector scaled so that a resultant vector of length 1 (if all dipoles were strictly collinear and pointing in the same direction) would span the whole radius of the plot. **d. Average projection of the dipoles in sensor space**. Black circles identify sensor positions on the topographical view; the nose is at the top of the view, and the left side of the head appears on the left. The color bar indicates the strength of magnetic field exiting (red) and entering (blue) the head in femtoTeslas (fT). **e. Power contour plots for tests on simple amplitude differences at this location**. Color represents the statistical power estimated by Monte Carlo simulations, i.e. the number of significant tests divided by the number of simulations (500) for all tested combinations of trial and subject numbers. Black isocontour lines on the plots highlight spline-interpolated power estimates of 0.5 and 0.8. **f. Power contour plots for tests on differences of squared amplitudes at this location**. The same conventions as in e apply. **g. Power contour plots for tests on differences in GFP at this location**. The same conventions as in e apply.

Unlike the fusiform source, the source in the superior occipital gyrus, had a much larger variability in position across the 89 HCP subjects. This spatial variability is visible in Figures 7a and 7b. We note that, although relatively limited compared to some other sources, orientation variability at this location was also larger than in the fusiform cortex (Figure 7c). Together, these differences yielded a relatively lower amplitude of signal at sensor level (Figure 7d), as compared to the previous dipole in the fusiform gyrus. Accordingly, the detectability of this source was poor, with no tested number of subjects and trials reaching 50% statistical power (Figure 7e). When using the difference of squared amplitudes at sensors, or of GFP, to compare conditions statistical power was generally increased.

**Figure 7.**
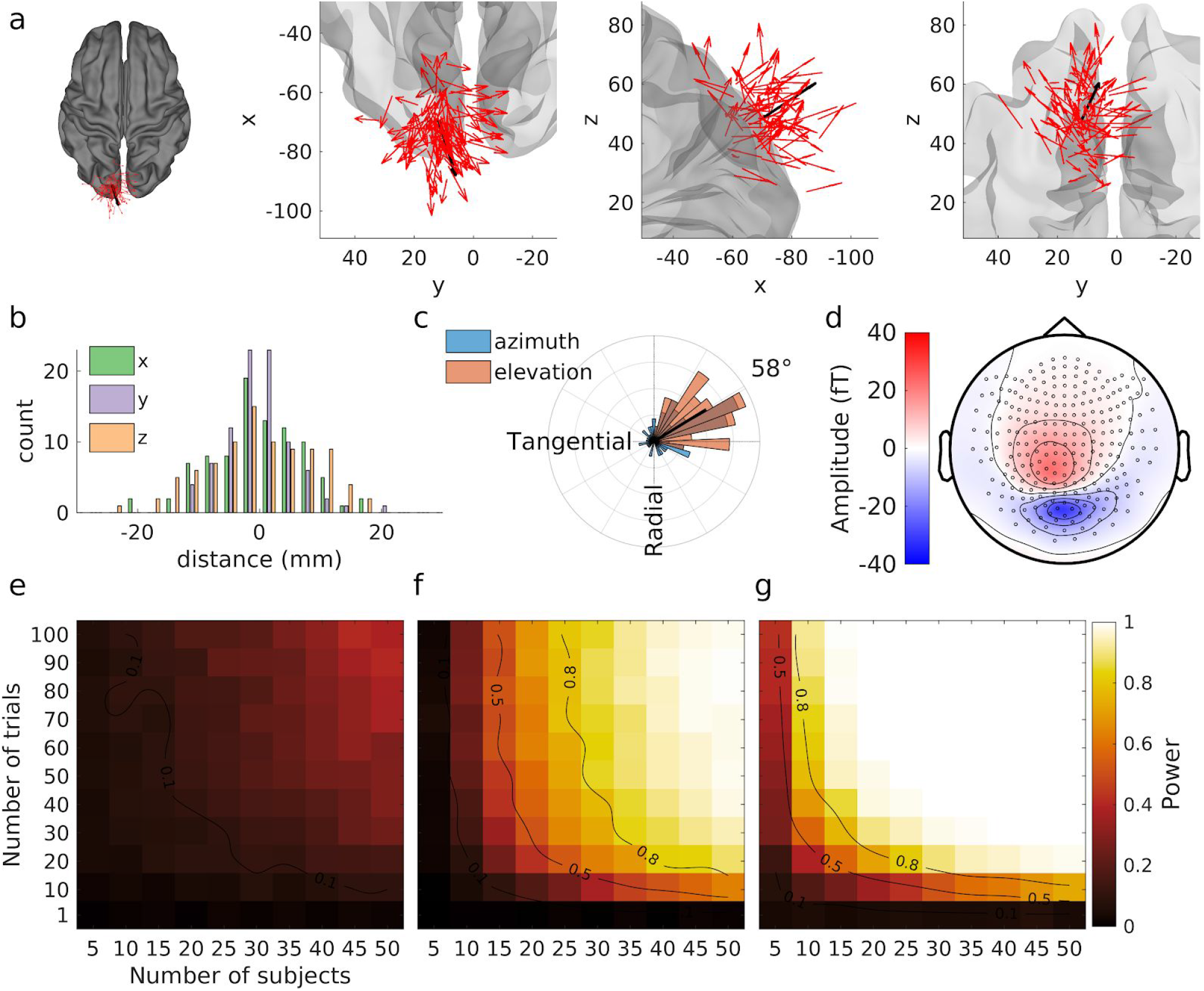
Detecting sensor-level effects for a source with high cross-subject position variability in the superior occipital gyrus. The same conventions and legends as for Figure 6 apply. See main text for further explanation.

The two dipole locations presented above illustrate qualitatively the effect of position variability on detectability. However, as already mentioned, co-variation in the different spatial properties of anatomically constrained sources is a confound that undermines robust conclusions specific to position variability. Therefore, we moved to a more specific manipulation of position variability, maintaining other variables constant, in simulations unconstrained by anatomy. Here, we selectively allowed dipoles to change position across subjects by sampling positions at random from a 3D normal distribution with a set standard deviation across subjects. Figure 8 shows the effect of this manipulation on statistical power for five equally spaced position variabilities ranging from 0 to 10 mm standard deviation. For all explored position variabilities, the initial dipole position was selected on the anterior bank of the precentral gyrus, hence average position of dipoles across participants tended toward that same position (Figure 8a). The projected signal at sensor level is illustrated in Figure 8b, showing little effect on signal amplitude, and in turn little effect on estimated statistical power (Figure 8c).

**Figure 8.**
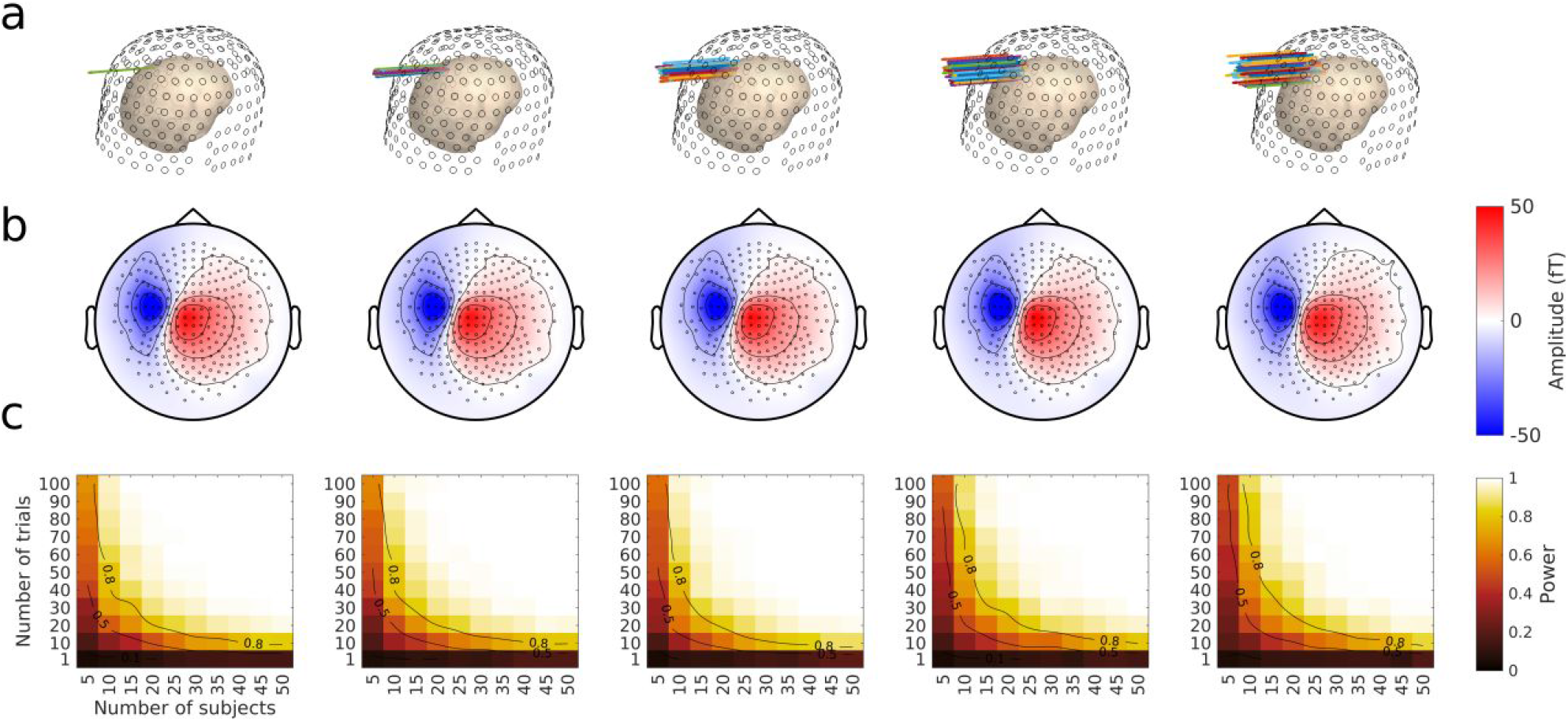
Detecting sources with varying cross-subject position variability. **a. Locations of individual dipoles**. Source dipoles in 89 subjects (colored arrows) are represented on an individual head model (pinkish inner surface) relative to sensor positions (black open circles on outer surface). The source dipoles spanned one of five linearly increasing standard deviation values around the initial source in the precentral gyrus (see main text), ranging from 0 (no variability at all), to 1 cm standard deviation in radius. **b. Average projection of the dipoles in sensor space.** The same conventions as in Figure 6d and 7d apply, however, the scale on the amplitude axis has been altered. **c. Power contour plots for each simulated level of source position variability.** These plots were obtained by Monte Carlo simulations as explained in Method and Figure 6e. The same conventions as in Figure 6e and 7e apply.

##### Interim Summary

Manipulating position variability had little effect on detectability when the manipulation was selective. In realistic settings, regions with high position variability across subjects such as the occipital cortex tend to be also more variable in terms of orientation. We explore further the effect of orientation variability across subjects in the next section.

#### Orientation variability across subjects

Figure 9a shows cross-subject variability in orientation for every source location on the cortical surface, plotted on the average brain of the 89 subjects in the HCP dataset. The histogram of these vector lengths across all vertices is plotted in Figure 9b, which highlights the heterogeneity of orientation variability across the brain. Overall, values ranged between 0.0019 and 0.74, with maximal values in lateral occipital regions, and minimal values along the central sulcus, the ventral prefrontal cortex, dorsomedial prefrontal cortex and insular cortex. Dipoles in the most variable regions of the lateral occipital cortex could be oriented up to 180° apart (i.e. in opposite directions) across subjects, dramatically reducing the net contribution to the signal at sensor level across a group of subjects.

**Figure 9.**
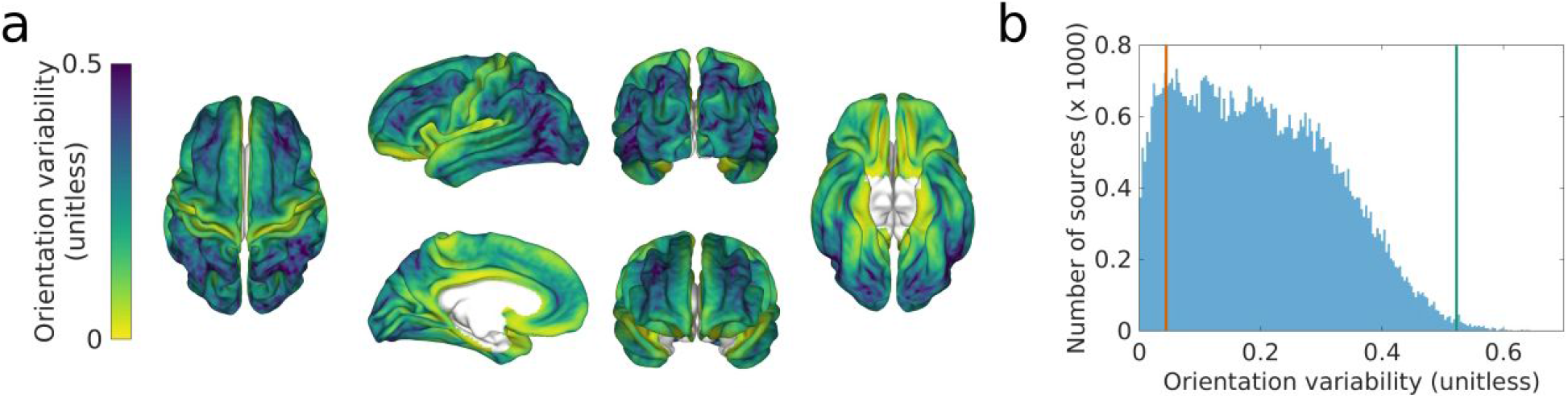
**Average cross-subject variability in orientation** shown on the average brain of 89 subjects in the HCP data. **a. The cross-subject variability in orientation** is the log of the inverse of the average resultant vector length across individual sources, for each cortical location. It is shown on superior, left lateral, left medial, posterior, anterior, and inferior views of the brain (from left to right and top to bottom). **b. Histogram of the distribution of cross-subject variability across all cortical vertices.** The red and green vertical lines indicate the orientation of the two dipoles that we selected to illustrate the effect of this parameter in Figures 10 and 11, respectively.

Orientation variability is expected to have an impact on detectability because dipoles with different orientations, even if they share the same position, project a magnetic field that has a different topography at the sensor level. Contrary to position variability, changes in orientation can change the polarity of the magnetic field at a given sensor even with minimal orientation change. Hence a major effect on statistical power may be expected. Here, we review two example locations, one with relatively low orientation variability across subjects, in the insula (x=6, y=36, z=30), and the second one with large variability, in the posterior superior temporal sulcus (x=−47, y=42, z=30).

For the insula source, Figure 10a shows that most individual sources are parallel (see in particular the middle zoomed inset), resulting in a length of the resultant vector of 0.90, shown as a black line in Figure 10c, and an average projection to sensors with relatively high amplitude as can been in FIgure 10d. It may be noted that the position variability of this source was also limited (Figure 10b). The power to detect this signal is such that 20 subjects with 90 trials each, or 50 subjects with 30 trials each allow detecting the 10 nA.m source with 80% statistical power (Figure 10e). When using the difference of squared amplitudes at sensors, or GFP, to compare conditions statistical power was generally decreased.

**Figure 10.**
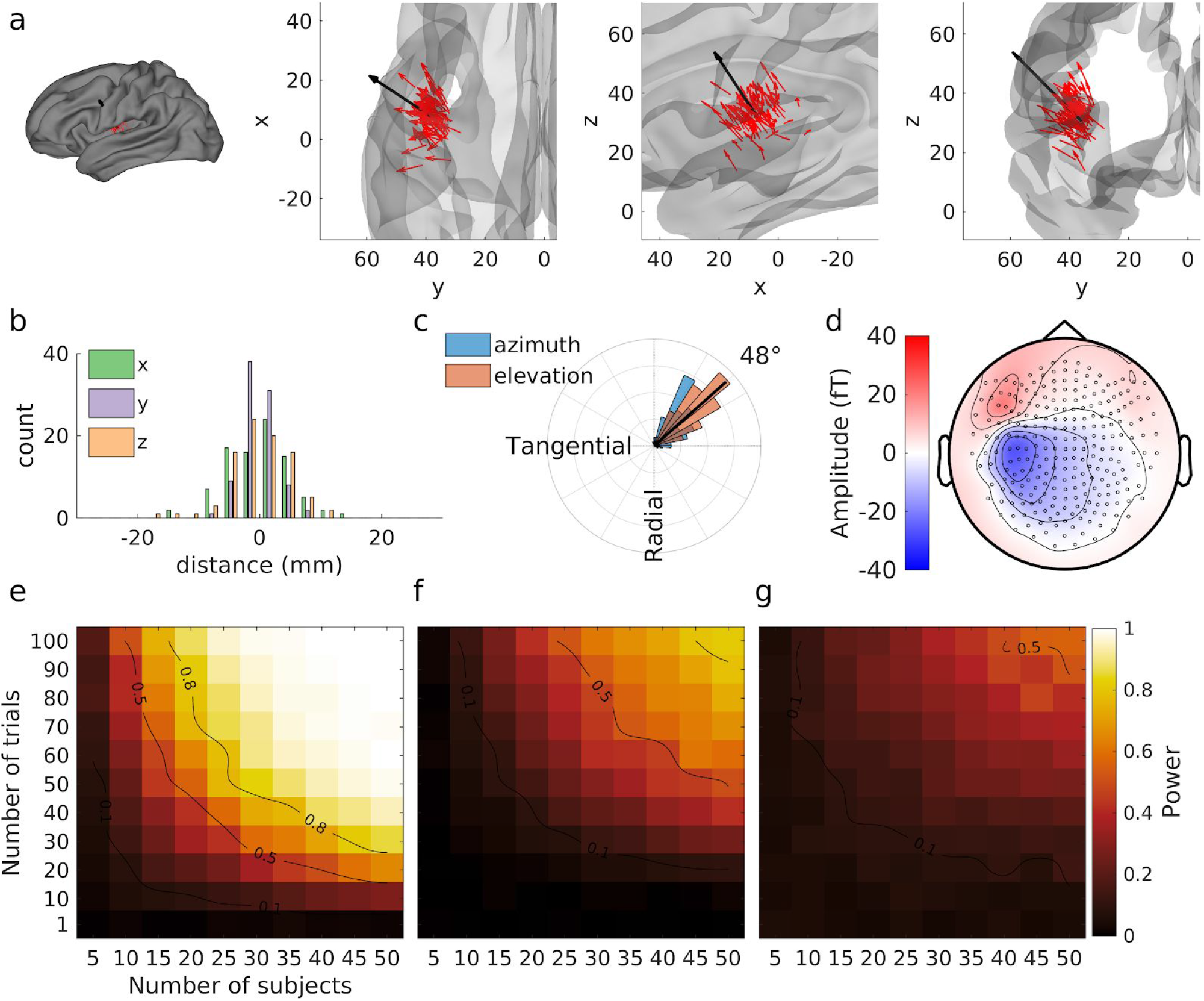
Detecting sensor-level effects for a source with low cross-subject orientation variability in the insula. The same conventions as for Figure 6 apply. See main text for further explanation.

The source in the posterior superior temporal sulcus (pSTS) on the other hand showed a highly variable orientation across subjects (Figure 11a and c), with a resultant vector length of 0.30. We note that although we tried to select a source with high orientation variability while keeping its other spatial properties similar to the previous example source, the position variability of the pSTS source was relatively important. This source resulted in a very weak average amplitude of the signal projected at sensor level (Figure 11d). Accordingly, this signal was virtually undetectable within the range of explored subjects and trial numbers (Figure 11e). When using the difference of squared amplitudes at sensors, or GFP, to compare conditions statistical power was generally increased.

**Figure 11.**
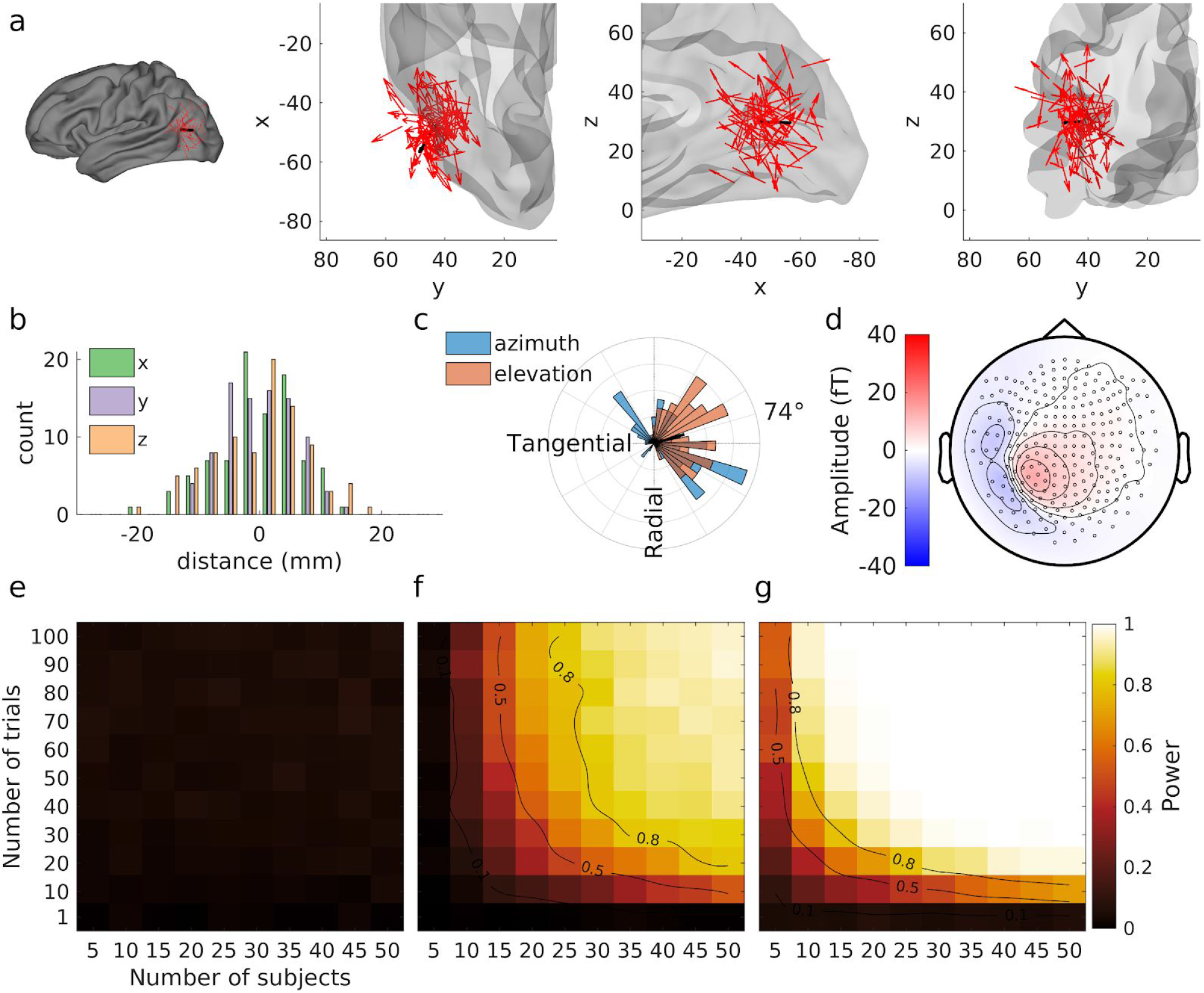
Detecting sensor-level effects for a source with high cross-subject orientation variability in the posterior superior temporal sulcus. The same conventions as for Figure 6 apply. See main text for further explanation.

The two dipole locations presented above illustrate qualitatively the effect of orientation variability on detectability. To examine this effect more systematically, we next move to a specific manipulation of orientation variability, in simulations unconstrained by anatomy. This time, position was held constant and only orientation was varied across subjects. We sampled orientations at random, adding a normally distributed random azimuth and elevation to the original orientation (arbitrarily set to the tangential source in the precentral sulcus examined in Supplementary Figure 6), with a standard deviation across subjects set to five evenly spaced values between 0° (fixed orientation) and 180°. The effect of this manipulation on statistical power is illustrated in Figure 12. It is noteworthy that the effect of orientation variability is most visible for random orientations with a standard deviation above 90°. This is arguably due to the fact that dipoles cancelling each other out (i.e. oriented in opposite directions) can only occur when the distribution of orientations is sufficiently broad.

**Figure 12.**
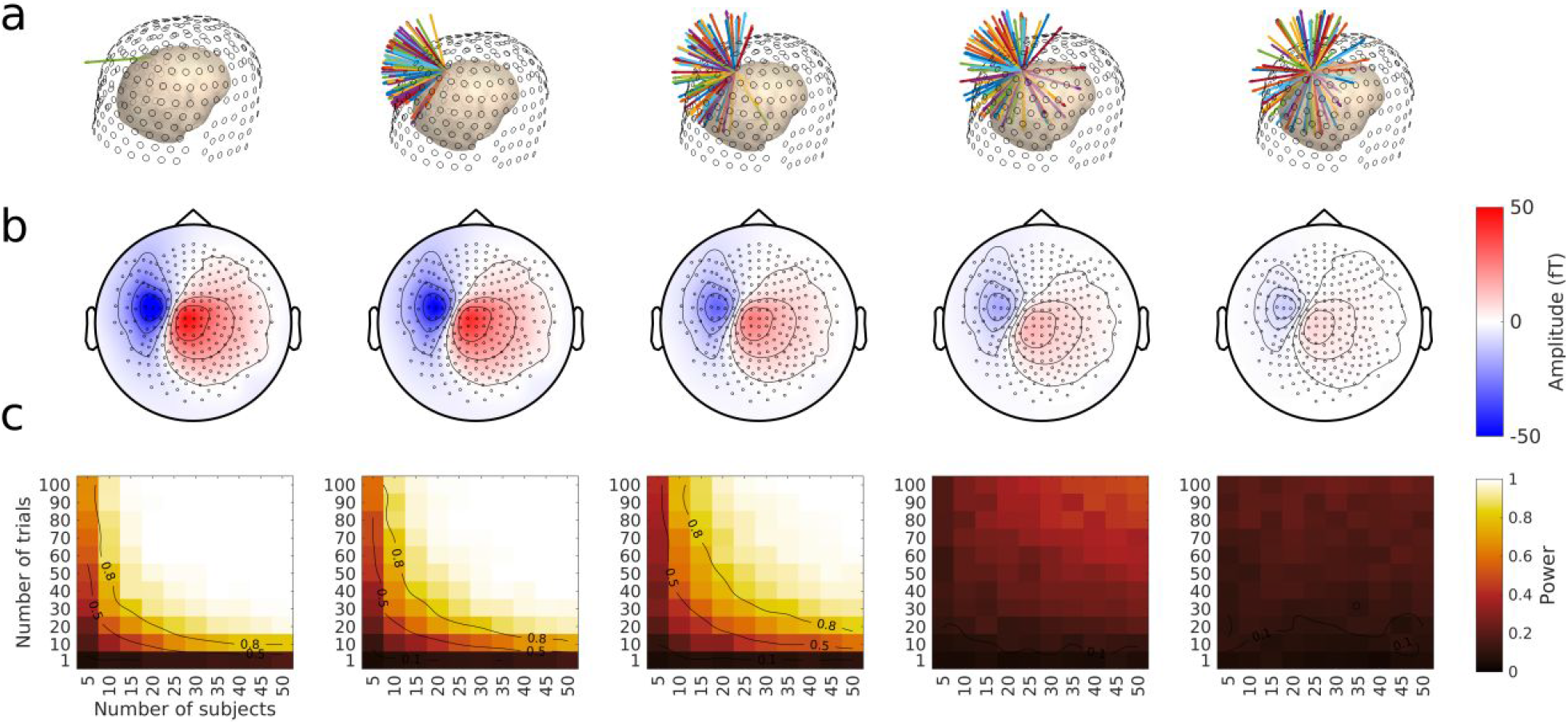
Detecting sources with varying cross-subject orientation variability. The same conventions as in Figure 8 apply. See main text for further explanations.

##### Interim Summary

We showed that second-level properties of the sources have very different effects on detectability. Within the range of variabilities explored, orientation variability had a much larger effect than position variability. Together with the effect of the first-level properties examined above, these results highlight the importance of considering source properties when planning a well-powered MEG experiment. We discuss this and other issues in more detail below.

## Discussion

Adequately powered MEG (and EEG) experiments require an appropriate sample size. In this study, we showed that the properties of the brain sources expected to contribute effects in a given experiment critically affect attainable statistical power in MEG. Specifically, we focused on the spatial properties of the sources and on their variability across subjects to examine how these factors affect group-level statistical power with classical tests at sensor level. We showed—quite expectedly—that the distance of the sources to the sensor array has a strong effect on detectability, with deeper sources being virtually undetectable with reasonable sample sizes for a single-session MEG experiment. We also observed another well-known effect, namely that sources oriented radially are almost undetectable in MEG data, as compared to other orientations. Most interestingly, we also found effects of source variability in location and orientation across subjects, which were not trivial and may also run counter to thinking about the detectability of neural sources. Finally, the type of measure (i.e. amplitude, squared amplitude, or GFP) used to test for differences also had a notable effect.

Examining the influence of cross-subject variability, we observed that source position variability across subjects had, in fact, little effect on statistical power, whereas orientation variability (or the lack thereof) strongly affected statistical power. Signal detectability at sensor level thus depends not only on the source origin (position and orientation), but also on the less well predictable cross-subject variability at the source origin. Therefore, one take home message for this study is that there is no simple solution for finding the optimal number of trials and subjects for all types of evoked MEG (or EEG) studies. Considering our present results, finding the optimal amount of trials and subjects for all types of MEG and EEG evoked studies would require considerable further methodological developments. One potential avenue could be to use Sarvas’s formula (Sarvas, 1987) as a starting point and analytically derive the effects of the properties shown here on statistical power. However, there will likely be variations in power across different brain structures within a single experiment that we believe will make calculations of power challenging. Thus, we emphasize the importance of considering the expected brain sources of activity, their anatomical location, and their cross-subject variability while planning studies in cognitive, social and systems neuroscience.

Resources such as the present paper (and the code distributed with it) could be used to predict the required number of subjects for detecting signals coming from a specific region. This should, however, be approached with some caution. The present work focuses on the spatial properties of the underlying neural sources. We did not attempt to model temporal variability - another important dimension in measuring functional activation. Furthermore, the only form of within-subject variability included in our simulations was in the sampled resting state data. In a real experimental setting, it would be important to take into account the fact that brain responses may never repeat twice in exactly the same way. Latencies and spatial locations are likely to be different from one trial repetition to the next in any given experimental condition. Moreover, properties such as source strength may compensate for some effects of spatial parameters (e.g., the effect of depth on source detectability may be compensated by source strength). Thus, all relevant parameters should be considered, when trying to predict statistical power for given sources in the real brain. We explicitly ignored these types of variability here in order to keep the problem tractable. Below we discuss some issues and considerations for planning MEEG studies, while keeping these limitations in mind.

Previous studies have explored the effects of spatial properties from a single-subject signal detection perspective, aiming to provide estimates of how much MEG signal can be detected given these properties and the amount of noise in an individual subject’s data (Ahlfors, Han, Belliveau, et al., 2010; Goldenholz et al., 2009; Hillebrand & Barnes, 2002). It was demonstrated that source depth relative to sensors is the most important factor affecting source localizability, with additional effects due to source orientation (Hillebrand & Barnes, 2002). Additionally, others have shown that MEG signal-to-noise varies considerably with source location (Goldenholz et al., 2009) and orientation (Ahlfors, Han, Belliveau, et al., 2010) in the brain. Pushing the analysis one step further, our group-level statistics approach takes a pragmatic stance on this issue, speaking directly to experimenters for whom changes in signal-to-noise ratio are particularly relevant for deciding on sample size, i.e. how many subjects or trials to include in an experiment.

It is worthy to note here that although the general reasoning of this study could be directly applied in EEG as well as MEG data, the effects of source orientation explored here cannot be directly applied to EEG, since the EEG field does not suffer from cancellation for radial sources, and the generally broader topographies elicited in EEG are also less prone to cancellation due to orientation variability.

Our study has identified challenges for estimation of statistical power in simulations of brain activity. Not only are there interdependencies between source parameters, but differences also arise due to the chosen brain activity measure (amplitude, squared amplitude, and GFP). As a consequence, we showed here that the source detectability question cannot be answered definitively. First, the large effects of spatial properties across cortical brain structures on detectability that we observed (Figure 2) are a major challenge for predicting statistical power. Indeed, not only do the parameters that we examine greatly vary across the brain (as shown by Figures 3, 4, 5 and 9), but also these parameters are not independent of one another across the cortical mantle (Supplementary Figure 2). For instance, sources with the shortest distances to sensors also tend to have a radial orientation, as gyral crests are generally found on the outer surface of the brain. Other regions with mostly tangential orientations, such as along the central sulcus, can be spatially highly consistent across subjects. In contrast, some brain regions have high degrees of inter-individual variability due to variations in cortical folding (e.g., pSTS, MT+/V5, inferior parietal cortex; Caspers et al., 2006), high degrees of curvature (e.g., occipital pole), or hemispheric asymmetries (Croxson et al., 2018; Ochiai et al., 2004). In addition, some deep brain structures, such as the hippocampus, may have less structural variability across subjects—this could mitigate some depth issues.

The choice of signal measure also influenced the ability to detect effects at the sensor level. Simple amplitudes seem to allow better detection of deeper structures, while squaring the data, or computing the GFP before contrasting conditions enhances statistical power for more superficial sources (Figure 2). Determining the parameters that govern how these transformations affect statistical power was beyond the scope of the present paper and will be explored in future simulation studies. Altogether, variations of power across the brain for a given sample size are difficult to predict in practice.

On a methodological note, we come to a forked path. Is it better to perform a realistic simulation without being able to fully disentangle the source of variations, and eventually obtain a more practical power estimate e.g. needed for planning new studies, grants etc.? Or, is it better to simulate unconstrained by anatomy, to systematically explore each parameter without considering interdepencies between source parameters? We believe that there is an important role for both types of simulations—to generate a better understanding of what is being measured, so that analysis methods can be improved in the future. In the present case, using these two types of simulation has shed light on some unexpected effects on signal detectability. Specifically, we have observed unpredicted enhancements or decreases in detectability that we would not have observed with tightly-controlled anatomically unconstrained simulations.

Statistical inference is usually made at the group-level. Predicting the adequate number of subjects for a given study also implies considering between-subject variability in different cortical regions. Although first-level spatial properties and their effect on signal are straightforward to measure and model with Maxwell’s equations, second level properties and their effects on group-level statistics are much more challenging to model. Our approach was to directly manipulate anatomical variations in our simulations free from anatomical constraints. Our first observation was that strictly parallel dipoles tend to combine across subjects, even when their position is relatively spread out, thus creating a more readily detectable net magnetic field. Randomly oriented dipoles on the other hand tend to cancel each other out. Beyond a certain orientation variability, statistical power becomes critically low, giving no opportunity to detect a signal even with a large number of subjects. This observation has an interesting implication: when comparing our whole brain power analysis for a sample of 25 subjects (Figure 2) with the whole brain signal-to-noise mapping for a single subject in MEG obtained by Goldenholz et al. (Goldenholz et al., 2009, their Figure 2, first row), at least two differences are apparent: First, some regions with high SNR e.g. in the occipital/parietal cortices, remain hard to detect at the group-level, at least if one relies on the amplitude difference. Second, other regions with low SNR, in particular on the medial and ventral surfaces of the brain, are readily detectable at the group level (e.g. in anterior cingulate cortex, or ventromedial prefrontal cortex). We hypothesize that cross-subject variability explains these discrepancies. On the one hand, it is thanks to a particularly consistent source orientation across subjects that some regions with low SNR in Goldenholz’ study still show high power in our study, and on the other hand other regions with high SNR are less detectable at the group level due to high variability in source orientations across subjects. In other words, the poor detectability of some cortical regions seems to be mitigated by their anatomical consistency across subjects (and conversely some highly detectable regions become less visible at group level). Most importantly, overall, we believe that there is no unequivocal answer to the question of an appropriate sample size in MEG (or EEG) experiments. Our study highlights the importance of considering between-subject variability in addition to the long-known spatial properties of sources when it comes to deciding how many subjects and trials to include in an experiment.

An important point to note regarding the decreased detectability with higher between-subject source variability is that it is standard procedure in MEG research to localize sources in individual subjects on their own anatomy before combining data for group analysis in source space (Hari & Puce, 2017; Jas et al., 2018). Going for source-level analysis at the individual subject level can avoid a large portion of the problem of between-subject variability, provided that data at the cortical level are properly aligned across subjects. Specifically, the cancellation effects we observe here across subjects when sources of different subjects point in opposing directions (Figure 11) should be largely avoided when sources are estimated in individual subjects first, then aligned across subjects on a template before being averaged (Hari & Puce, 2017). Further studies will have to examine the limits of this reasoning in detail. In particular, it will be important to examine group-level source detection in detail if we want to formulate more precise recommendations in the future.

As mentioned in introduction, a group-level approach has recently been used by Boudewyn et al. (2018) for EEG data in sensor space, exploring the dependency of statistical power on sample size (subjects and trials), while focusing on effect size at the electrode level for a set of ERP components. Our approach here is similar, but instead of starting from an expected ERP difference at sensor level, we started from the expected neural source. This complementary approach may allow principled investigations beyond known ERP effects, and could allow a better planning of studies targeted at activating specific brain regions. It also has potentially strong practical implications as power analyses are usually mandatory in grant applications, or for preregistering studies.

Another important aspect to consider for future studies will be that of spatially extended sources. Here, we only modelled dipoles at single vertices—a limitation of our study. Although dipolar sources are easy to model and often used to appreciate detectability, more realistic extended patches of activity can have a difficult-to-predict net effect at sensors. For instance, such extended patches that occur in the homologous regions of the two hemispheres on the mesial wall, or that are spread across opposite walls of a sulcus, can show relatively lower detectability due to opposite sources cancelling each other out (Ahlfors, Han, Lin, et al., 2010; Fuchs et al., 2017; Hari & Puce, 2017). It should be noted, however, that functional area borders often tend to follow gyral and sulcal crests, making synchronous activation on both sides of a sulcus less likely (Destrieux et al., 2010; Glasser et al., 2016). The detectability of extended sources at the group-level in such cortical regions needs to be explored in the future for additional practical recommendations.

High-temporal resolution is desirable in clinical and research studies of cognitive and social neuroscience. This is particularly true where the precise detection of timely neural activity is key, such as in detecting sources of epileptogenic spikes and seizures, as well as in hyperscanning and naturalistic protocols. It will therefore be of importance to extend explorations of statistical power to the time domain, in order to characterize how it may change over time, particularly before and immediately after stimulus onset, and examine effects on group-level signal detectability at specific latencies. Moreover, we think that investigations in sensor space will remain important in the foreseeable future, in addition to those in source space, because portable EEG and room temperature MEG studies are more likely to use relatively low numbers of sensors, making it potentially unrealistic to transform these data into source space.

Given the current emphasis on whole-brain data collection in cognitive, social and systems neuroscience, the explosion of network science based data analyses (Bassett & Sporns, 2017), and the modulation of activation as a function of perceptual and cognitive manipulations (Medaglia et al., 2015), it is important to understand how statistical power can vary across the brain and the critical dimensions along which these variations occur. Critically, the statistical power for detecting activity across a given network will *at most be only as good as its least detectable node*. It is thus important while planning an experiment to consider sample size in regard with the set of brain regions believed to be involved, especially when envisioning a network analysis. We hope that the present paper helps raising awareness about this and provides helpful information to the researchers for deciding on trial and subject numbers while balancing on experiment duration and budget constraints.

## Supporting information

Supplementary results

## Acknowledgements

Maximilien Chaumon and Nathalie George work on the CENIR MEG-EEG platform at the Paris Brain Institute, which has received infrastructure funding from the programs “*Investissements d’avenir*” ANR-10-IAIHU-06 and ANR-11-INBS-0006. Aina Puce received support from the College of the Arts & Sciences at Indiana University, Bloomington (Sabbatical Leave).

## Footnotes

1 Described here: http://www.fieldtriptoolbox.org/faq/how_are_the_different_head_and_mri_coordinate_systems_defined/#details-of-the-4dbti-coordinate-system

