## Supplementary results for "Statistical power: implications for planning MEG studies"

#### Choice of signal measure differentially affects statistical power throughout the brain

Supplementary Figure 1 below shows the difference in estimated statistical power between three types of signal measures. **a.** Amplitude vs. squared amplitude (depicting the difference between Figure 2a and Figure 2b in main text). **b.** Amplitude vs. GFP (depicting the difference between Figure 2a and Figure 2c in main text) **c.** Squared amplitudes vs. GFP (depicting the difference between Figure 2b, and Figure 2c in main text).

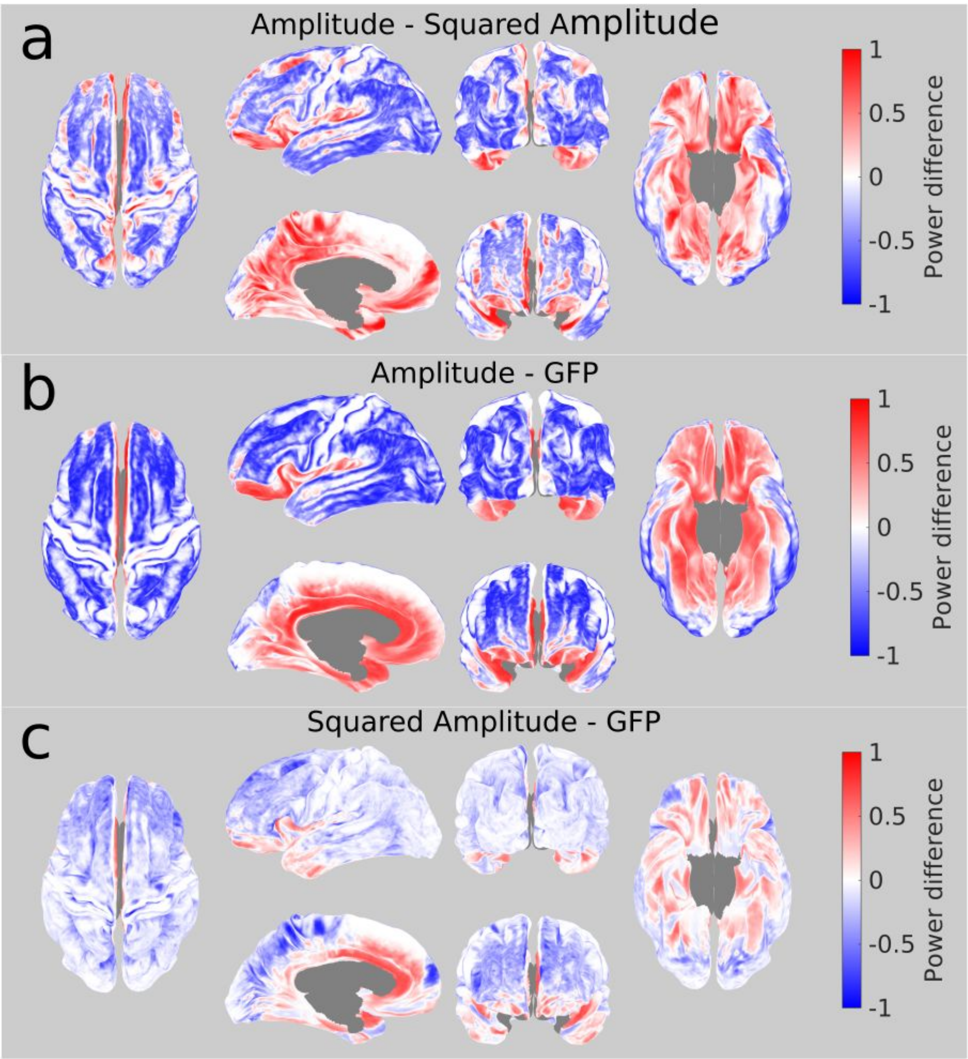

***Supplementary Figure 1: Difference in estimated statistical power across the cortical surface, depending on the chosen signal measure.*** *We considered the statistical power for detecting (at MEG sensor level) activity generated by a single dipole placed in turn at every possible cortical position, in simulated experiments with 25 subjects with 50 trials each. This activity was measured as signal amplitude, squared signal amplitude, or GFP, and statistical power, computed for each type of measure (as illustrated in Figure 2 in the main text). We then computed the difference in statistical power obtained for each type of measure. Images of superior, left lateral, left medial, anterior, posterior, and inferior brain views are depicted.* ***a.*** *Difference in statistical power between amplitude and squared amplitude measures.* ***b.*** *Difference in statistical power between amplitude and GFP measures.* ***c.*** *Difference in statistical power between squared amplitudes and GFP.*

#### Covariation among first- and second-level spatial properties

The spatial source properties used in the study strongly covary in a way that is not linearly determined. Characterizing these covariations is beyond the scope of this paper, but is illustrated in the figure below.

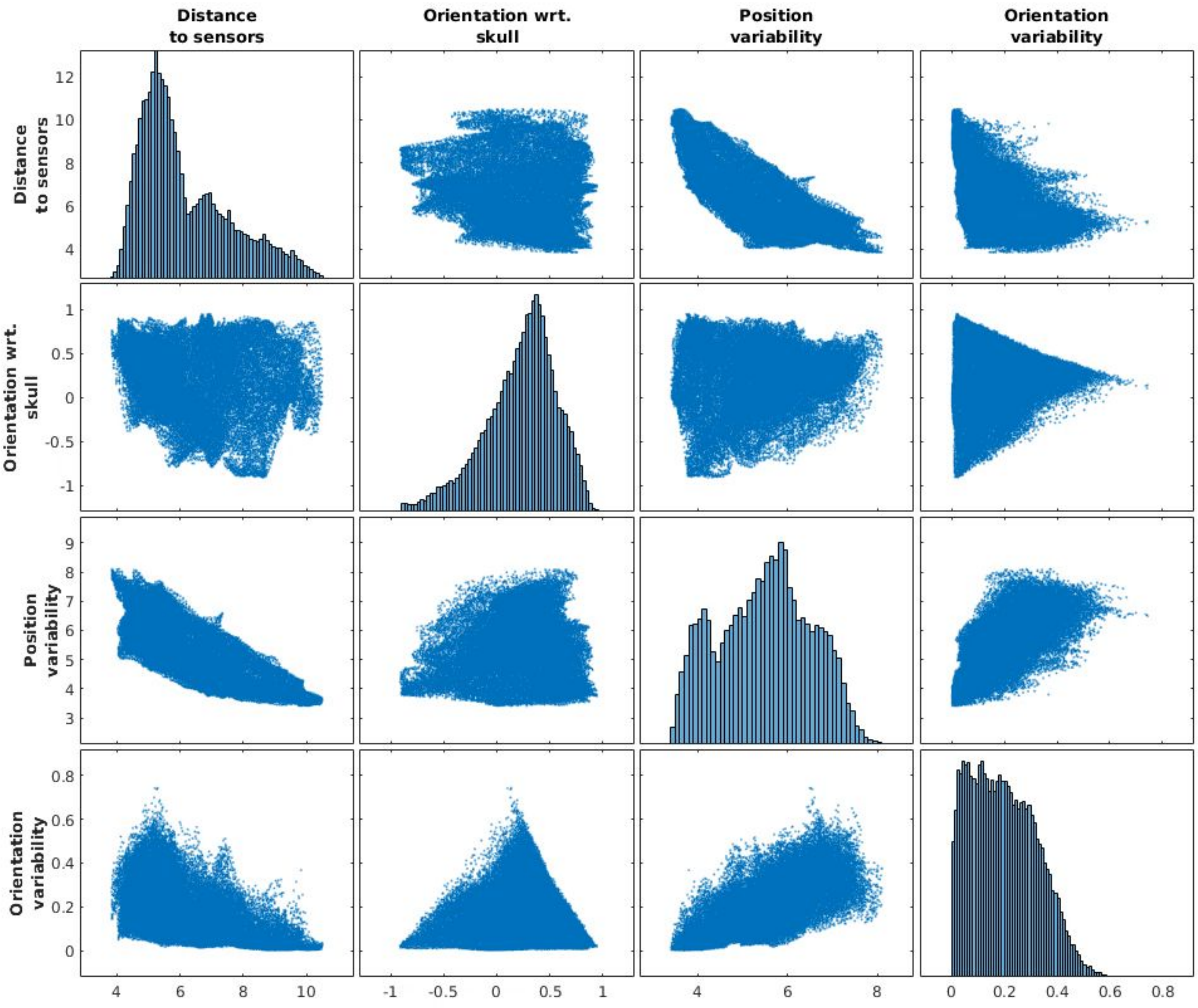


***Supplementary Figure 2: Covariation among spatial properties of sources across the brain.*** *The diagonal histograms show the distribution of each property (averaged across subjects), reproducing the insets in Figures 3, 4, 5, and 9 of the main text. The off-diagonal scatterplots show the covariation amongst every pair of variables. Note that the figure is symmetrical along its diagonal.*

#### Manipulating 'first-level' spatial source properties

The observed heterogeneity of statistical power across the brain [see Figure 2 main text] calls for a better understanding of the influence of source properties on detectable responses *at the sensor level*. In this Supplementary Results section, we display the results of simulations for first-level properties i.e. source position and orientation. We present two example locations in the brain to illustrate realistic variations in each property. These realistic variations are always concomitant with changes in other properties. We therefore also ran simulations for each source property unconstrained by brain anatomy.

### Distance

Based on physics, the distance to sensors is expected to have a large effect on signal amplitude, and thus on detectability of brain responses at sensor level (Hämäläinen, Hari, Ilmoniemi, Knuutila, & Lounasmaa, 1993; Hillebrand & Barnes, 2002; Malmivuo, Suihko, & Eskola, 1997). Here we illustrate how group-level statistical power changes with the numbers of subjects and trials for a 10 nA.m dipole placed in two locations—one in the middle frontal gyrus and the other in the orbitofrontal gyrus.

The first location in the middle frontal gyrus (x=71, y=21, z=41 mm) is at a distance of 5.6 cm to the closest sensor (on average across subjects). Supplementary Figure 3 presents simulation results for this dipole. An identical figure structure will be repeated for illustration of other source locations in the figures that follow. Supplementary Figure 3a displays the chosen dipoles across the 89 subjects of the HCP MEG dataset as well as the average dipole across all subjects. The position variability across subjects can be well seen in these top panels, as well as in Supplementary Figure 3b, which shows the histogram of dipole positions relative to the average position across subjects, indicating that the dipoles spread within +/-

10 mm from the average position in most subjects. Similarly, Supplementary Figure 3c shows the distribution of dipole orientations with respect to the closest point on the subjects' head across subjects. This plot indicates that the orientation for the chosen middle frontal gyrus dipole is on average 69° away from the radial orientation (i.e. away from the orientation that points exactly in the same direction as the closest sensor). Supplementary Figure 3d shows the average projection of this dipole to the sensor array, revealing a high signal amplitude at frontal sensors. This amplitude, when added to the resting state data could be detected with a standard paired t-test across subjects (corrected for multiple comparisons across sensors) against resting state data alone, with the estimated statistical power level shown in Supplementary Figure 3e for varying numbers of subjects and trials. In this particular case, achieving 80% power to detect the dipole signal requires at least 20 subjects with ~50 trials per subject, or up to 50 subjects with only 10 trials per subject. When using the difference of squared amplitudes at sensors to compare conditions, statistical power was decreased for all trial and subject numbers. When using the difference of GFP, statistical power was affected less dramatically in this case.

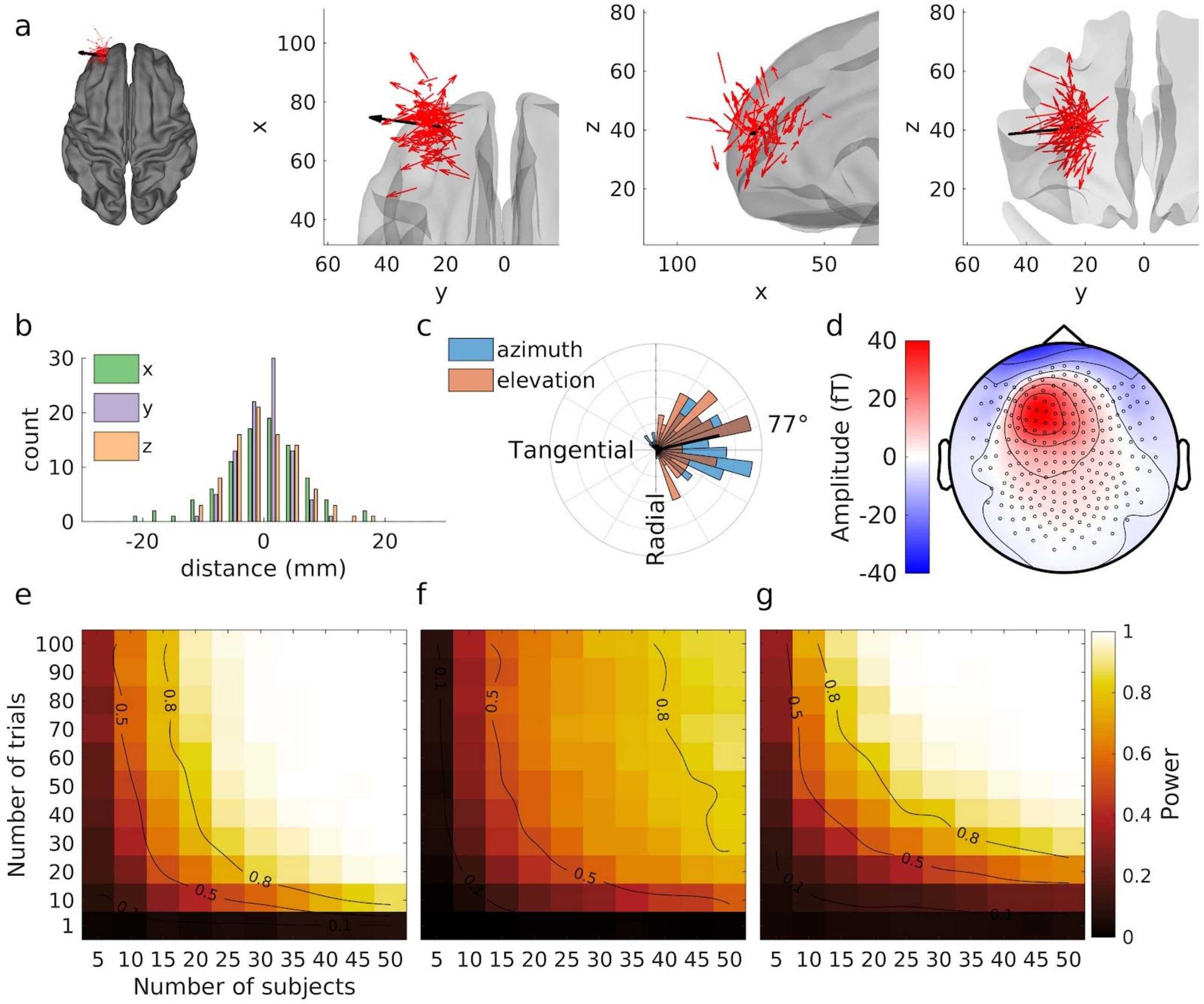


***Supplementary Figure 3. Detecting sensor-level effects for a source close to the sensors in the middle frontal gyrus. a.*** *positions of individual dipoles in 89 subjects (red arrows) are represented on the average brain in a superior overall view, and in zoomed superior, lateral, and posterior partial brain views. The bold black arrow in each panel shows the resultant vector or average dipole across 89 subjects.* ***b. Histogram of the distance of each dipole to the mean position of all dipoles in the three cartesian dimensions. c. Polar histogram of the orientations of the individual dipoles relative to the orientation of the closest point on the sphere encompassing the subjects' head.*** *Azimuth and elevation are referenced to the closest point on the sphere surface, so an orientation orthogonal to that of the sphere encompassing the subjects' head is shown as tangential (90°), and the colinear orientation is shown as radial (0°). The black thick line coming outwards from the center of the plot represents the orientation and length of the resultant vector scaled so that a resultant vector of length 1 (if all dipoles were strictly collinear and pointing in the same direction) would span the whole radius of the plot.* ***d. Average projection of the dipoles in sensor space.*** *Black circles identify sensor positions on the topographical view; the nose is at the top of the view, and the left side of the head appears on the left. The color bar indicates the strength of magnetic field exiting (red) and entering (blue) the head in femtoTeslas (fT).* ***e. Power contour plots for tests on simple amplitude differences.*** *Color represents the statistical power estimated by Monte Carlo simulations, i.e. the number of significant tests divided by the number of simulations (500) for all tested combinations of trial and subject numbers. Black isocontour lines on the plots highlight spline-interpolated power estimates of 0.5 and 0.8.* ***f. Power contour plots for tests on differences of squared amplitudes. The same***

***conventions as in e apply. g. Power contour plots for tests on differences in GFP.*** *The same conventions as in e apply.*

After examining the effects on statistical power of this relatively close-to-sensors dipole, we now turn to the detection of a deeper dipole, situated in the orbitofrontal gyrus (Suppl. Fig 4, x=44, y=13, z=7), at a distance of 8.4 cm from the closest sensor (on average across subjects). Compared to the previous location just described, in spite of its slightly more consistent location (Suppl. Fig 4b) and orientation (Suppl. Fig 4c) across subjects, and an almost tangential orientation (85° away from the radial orientation), the signal from this dipole projects with a much weaker amplitude to the sensor array (Suppl. Fig 4d). It is accordingly less well detected than the previous dipole (Suppl. Fig 4e). At least 25 subjects with 90 trials each, or 50 subjects with at least 40 trials each are necessary to detect this dipole with 80% statistical power. When squaring amplitudes before computing their difference to compare conditions, statistical power was generally decreased for all trial and subject numbers. When using the difference of GFP, statistical power was decreased even further.

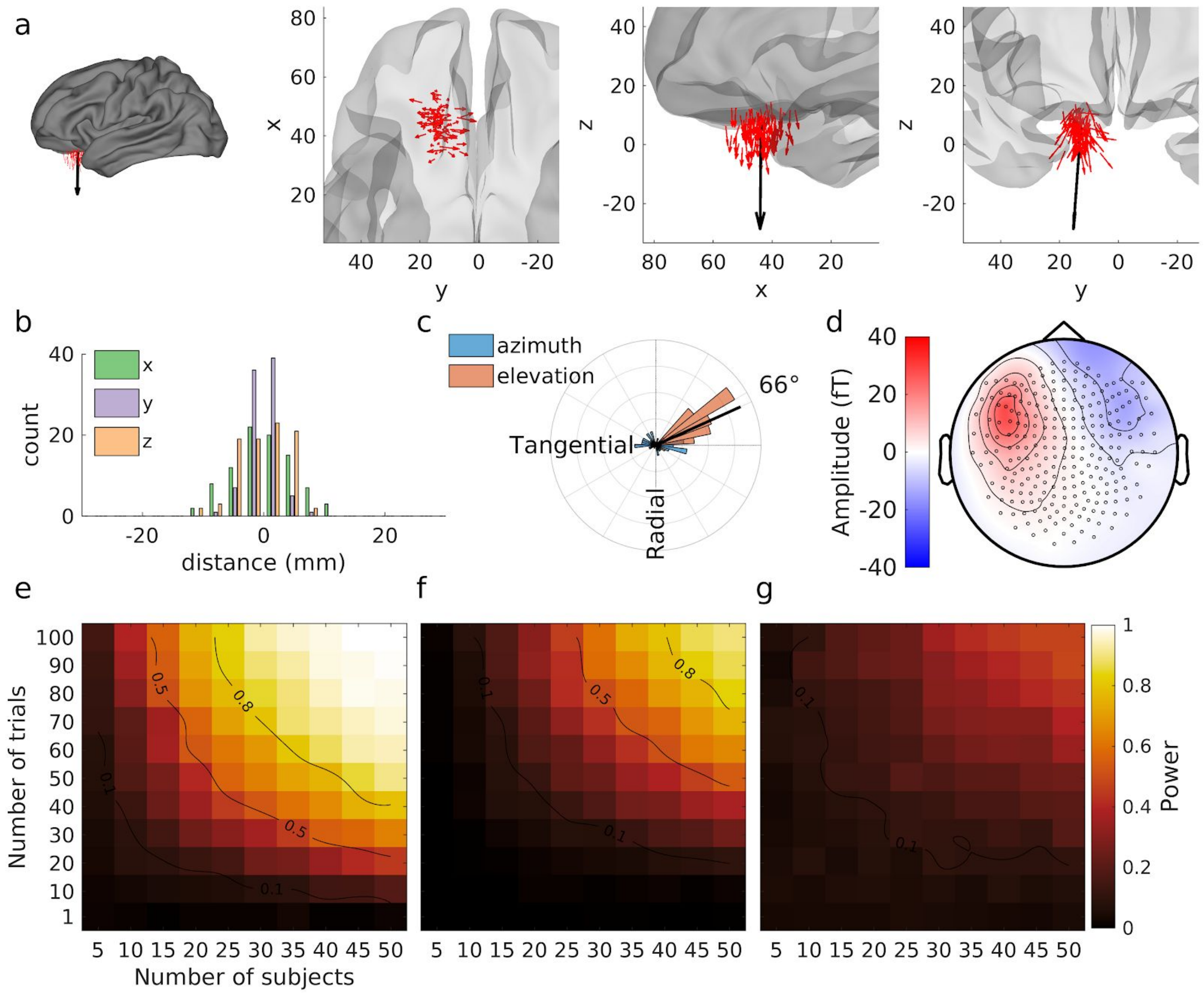


***Supplementary Figure 4: Detecting sensor-level effects for a deep source in orbitofrontal cortex.*** *The same conventions as for Supplementary Figure 3 apply. See supplementary text for further explanation.*

The two anterior frontal dipole locations presented above illustrate qualitatively the effect of distance to the sensors, but also highlight a clear limitation of this first approach of describing the effect of source properties on statistical power. Given how dipole properties covary in realistic source models across brain regions, it is nearly impossible to disentangle the respective contributions of distance, orientation, and their variability across subjects with these realistic source models. Therefore, we turned to a more selective manipulation of the distance to sensors by eliminating individual variation in other source properties—due to constraints of individual brain anatomies—in our simulations that appear below.

We selectively placed sources at set distances from sensors, along a radius running from an initial source position in the precentral sulcus (x=-12, y=33, z=70; a region close to sensors, detected with high power, as we later discuss; see Supplementary Figure 6) towards the center of the head, and ran simulations in the same way as before. Supplementary Figure 5 shows the effect of this manipulation on statistical power for distances to the closest sensor ranging from 4 to 12 cm. For all distances explored, dipoles were arranged in strictly the same orientation and relative positions (spatial variability in this case came only from the position variability of the initial source, Supplementary Figure 6a). The projected signal at sensors is illustrated in Supplementary Figure 5b, showing the sole effect of distance to sensors on signal amplitude. Finally, Supplementary Figure 5c shows how estimated statistical power varies across the distance parameter in these simulations. Note that our goal here is not to make a specific claim about statistical power at any specific distance, but rather to show how power varies with distance. In fact, we used dipoles with an amplitude of 5 nA.m in those simulations, i.e. half the amplitude used in the simulations constrained by anatomy, so as to not saturate completely the power plots. Under these conditions, our simulations show that distance to sensors greatly affects detectability: Sources that could be detected with 100% power for any number of trials above 10 in as few as 10 subjects when placed at 4 cm from the closest sensor were almost undetectable when placed at 10 cm away.

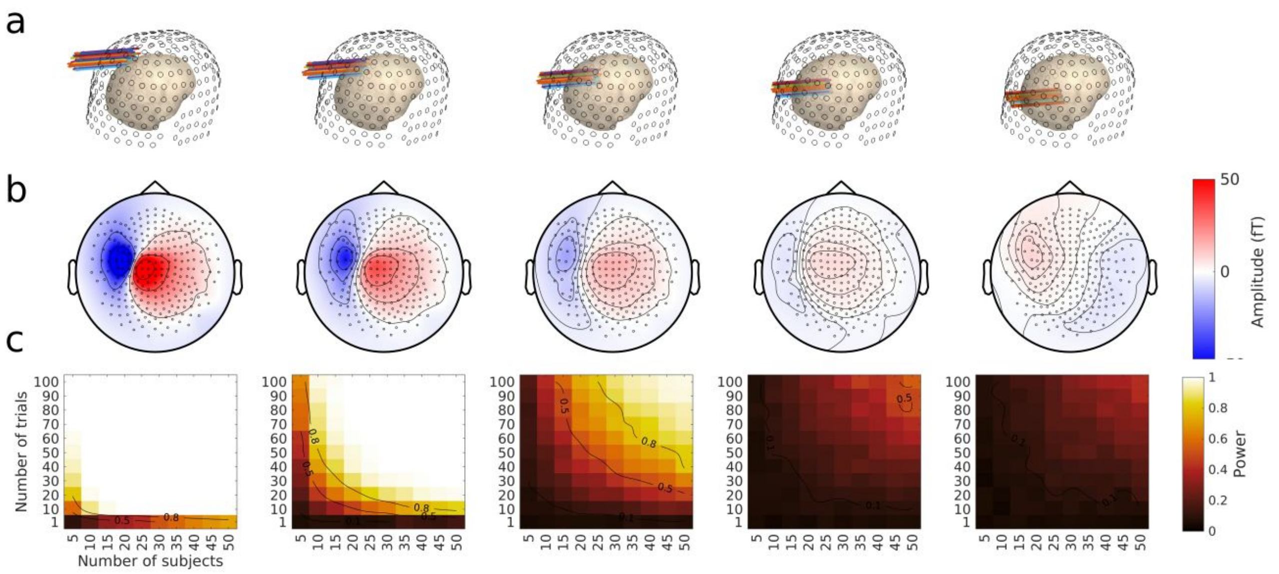


***Supplementary Figure 5. Detecting sensor-level effects for  sources at varying distances from the sensors. a. Locations of individual dipoles.*** *Source dipoles in 89 subjects (colored arrows) are represented on an individual head model (pinkish inner surface) relative to sensor positions (black open circles on outer surface). The source dipoles were placed at five equally spaced distances ranging from 40 to 120 mm from the sensor closest to an initial source in the precentral sulcus (see text), towards the center of the head (origin of the coordinate system).* ***b. Average projection of the dipoles in sensor space.*** *The same conventions as in Supplementary Figure 2d apply.* ***c. Power contour plots obtained by Monte Carlo simulations at each location, for amplitude measures.*** *The same conventions as in Supplementary Figure 3e apply.*

#### Orientation

Here, we use the same logic and display format as in the previous section. Based on physics, we expected the orientation of the sources relative to that of the head surface to have a large effect on signal amplitude (Hämäläinen et al., 1993; Hillebrand & Barnes, 2002; J. A. Malmivuo & Suihko, 2004), and thus on detectability.

We illustrate the effect of source orientation relative to sensors on group-level statistical power by examining two 10 nA.m dipoles at closeby locations in the precentral region, one in the posterior bank of the precentral sulcus, and the other on the gyral crest of the precentral gyrus . Each of these locations have very similar spatial properties, except for their orientation relative to the head surface. The sulcal source (x=-12, y=33, z=70) is close to a tangential orientation, while the gyral source (x=-4, y=37, z=78) is closer to a radial orientation. Our simulations show how the detectability of MEG signals at sensor level varies across numbers of subjects and trials for these two dipoles, as shown on the power plots of Supplementary Figures 6e and 7e. From these example sources, the effect of orientation appears to be dramatic, with 80% power for as little as 15 subjects and 40 trials, or 10 trials in 40 or more subjects for the first (more tangential) dipole, and no sufficient number of subjects and trials in our simulations to reach 80% power for the second (more radial) dipole. This illustrates the well-known effect of source orientation on MEG signal (Cohen & Hosaka, 1976). When squaring amplitudes prior to computing their difference, statistical power was decreased for all trial and subject numbers for the source close to tangential orientation, and increased for the source close to radial orientation. In contrast, for the source close to radial in orientation, power was generally greater. When computing GFP prior to computing the difference, statistical power was affected less dramatically for the source close to tangential orientation, but greatly increased for the source close to a radial orientation.

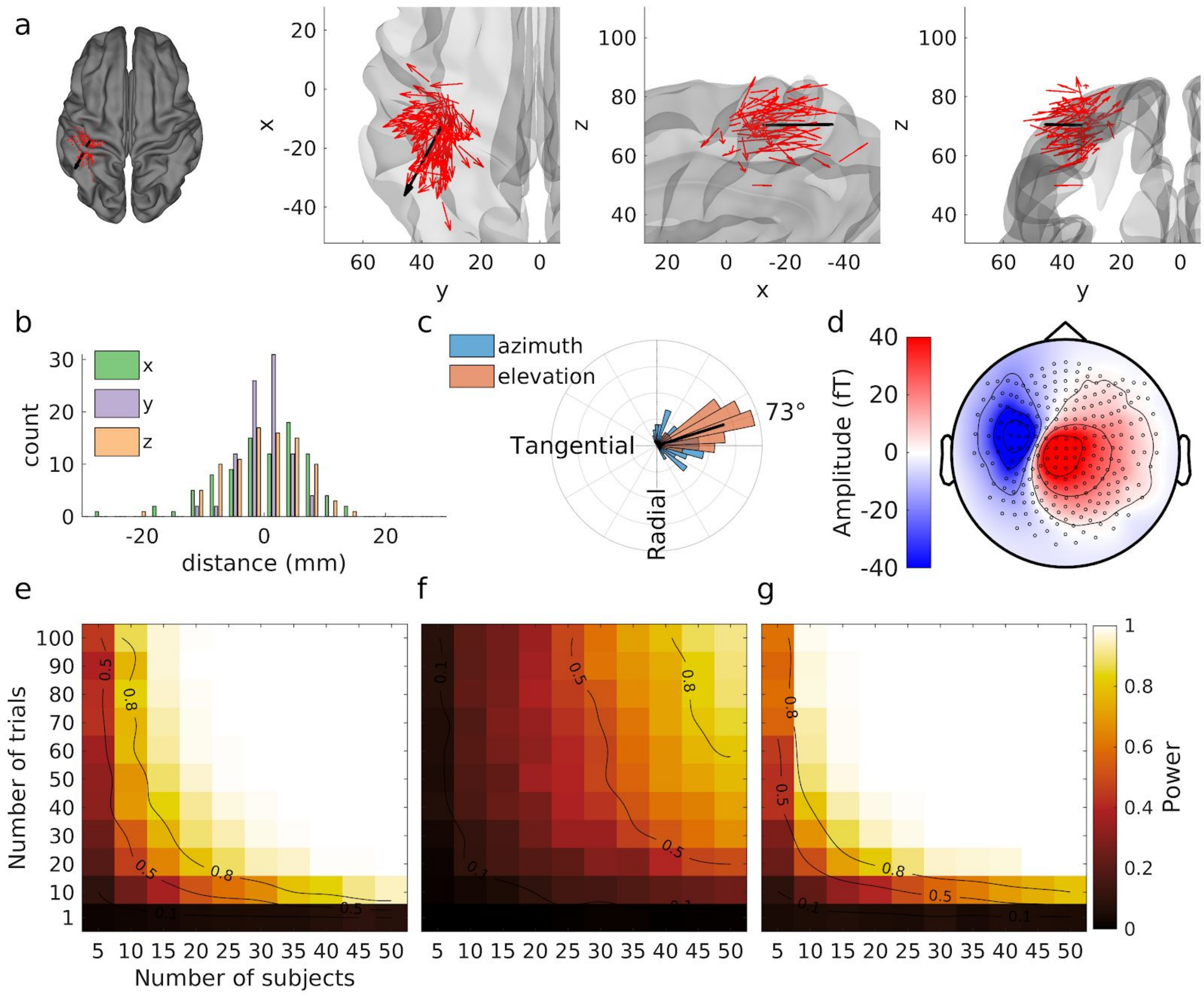


***Supplementary Figure 6. Detecting sensor-level effects for a tangential source in the precentral sulcus.*** *The same conventions as for Supplementary Figure 3 apply. See supplementary text for further explanation.*

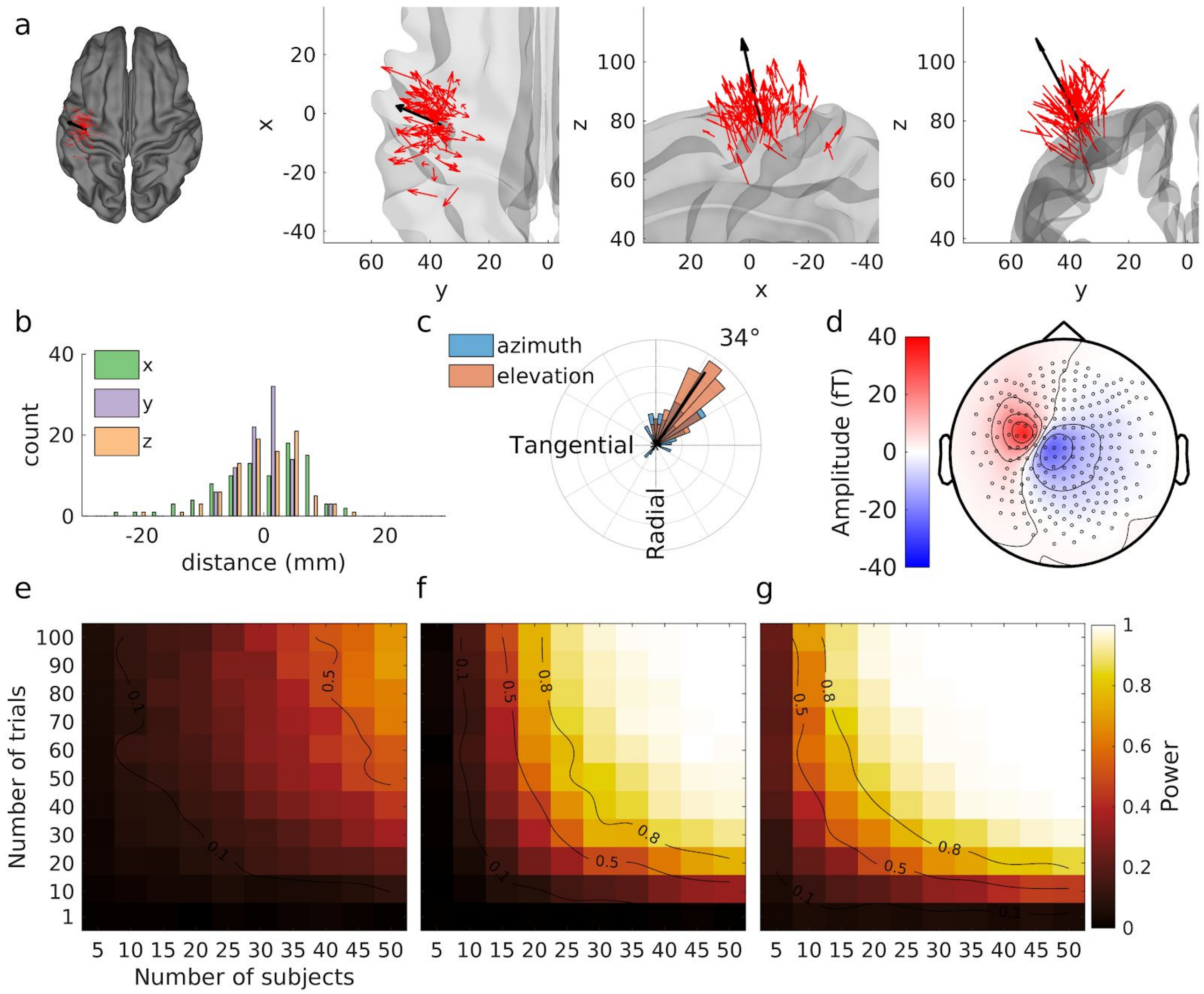


***Supplementary Figure 7. Detecting sensor-level effects for a radial source in the precentral gyrus.*** *The same conventions as for Supplementary Figure 3 apply. See supplementary text for further explanation.*

We now turn to a more selective manipulation of orientation by eliminating individual anatomical variation. Here, we chose a source at a set position and oriented it at set angles with respect to the closest point on a sphere encompassing the subject's head. We then ran simulations in the same way as previously. Supplementary Figure 8 shows the effect of this manipulation on statistical power for five equally spaced orientations ranging from 0° to 90°, starting from an original orientation strictly tangential to head surface, and progressively tipping around the y axis (passing through both ears), up to an orthogonal orientation (approximately tangential to the sphere encompassing the subjects' head; Supplementary Figure 8a). The projected signal at sensor level is illustrated in Supplementary Figure 8b, showing the effect of source orientation on signal amplitude. The most radial sources (left) produce a simulated signal that is about 10 times smaller than the most tangential sources (right), although with similar topography. Finally, Supplementary Figure 8c shows how estimated statistical power varied in these simulations from a situation where signal could never be detected above 40% with the number of subjects and trials explored for radial source orientations, to a situation where any number of trials above 10 to 20 in 15 or more subjects yielded 100% estimated power for tangential source orientations. Noteworthy, the decrease in detectability is narrowly focused at the radial orientation. A mere 22.5° shift away from this orientation (second orientation from the left in Supplementary Figure 8) produces a major improvement in detectability, with 80% power reached with for instance 50 trials in 20 subjects.

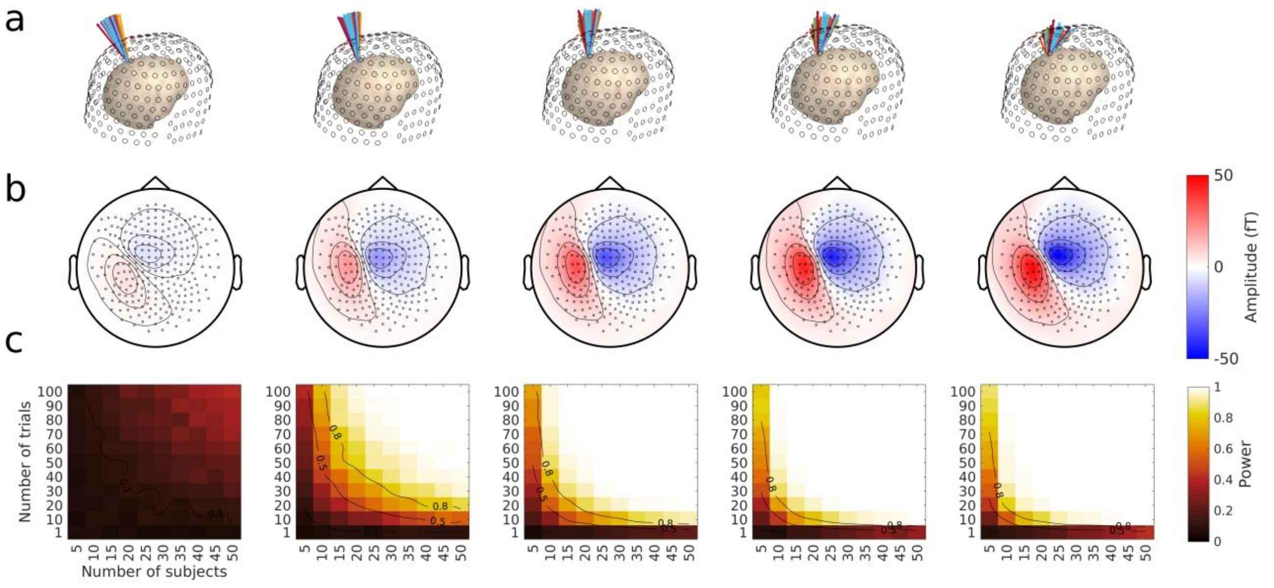


***Supplementary Figure 8. Detecting sensor-level effects for sources at varying orientations with respect to sensors.*** *The same conventions as in Supplementary Figure 3 apply. Orientations vary from 0°, i.e. radial orientation, on the left, to 90°, i.e. tangential orientation, in 5 equally spaced angles (0, 22.5, 45, 67.5, 90°). See supplementary text for further explanations.*

In summary, these supplementary results allowed us to explore the effect of the position and orientation of dipolar sources—our so-called "first-level properties". Quite predictably, we found that these first-level properties have a strong impact on signal detectability.
